# The proteome of remyelination is different from that of developmental myelination

**DOI:** 10.1101/2021.03.24.436006

**Authors:** Joana Paes de Faria, Maria M. Azevedo, Damaris Bausch-Fluck, Ana Seixas, Helena S. Domingues, Maria A. Monteiro, Patrick G.A. Pedrioli, Eduarda Lopes, Rui Fernandes, Chao Zhao, Robin J. M. Franklin, Bernd Wollscheid, Joao B. Relvas, Laura Montani

## Abstract

Loss of myelin underlies the pathology of several neurological disorders of diverse etiology. CNS remyelination by adult oligodendrocyte progenitor cells (OPCs) can occur but it differs from developmental myelination carried out by neonatal OPCs. We asked whether the myelin proteome of remyelinated regions is changed. We compared the myelin proteome formed during development to the remyelination proteome attained after lysolecithin-induced demyelination in the mouse spinal cord. Mass-spectrometry analysis of iTRAQ labelled myelin protein lysates showed that the proteome of remyelination is different from that of developmental myelination, leading to profound changes in myelin protein content. Aside from known mediators of oligodendrocyte differentiation, we found proteome alterations included modulators of metabolism, cell signaling and actin cytoskeleton dynamics. Downregulating one candidate (FSCN1/Fascin1) was sufficient to partially hamper oligodendrocytes *in-vitro*. In summary, we identify the difference in the proteome of remyelinating oligodendrocytes as a novel potential contributor to the pathophysiology of demyelinating disorders, thus providing new potential therapeutic targets for future studies.

## Introduction

In the CNS oligodendrocytes ensheath axons with myelin, a highly organized multi-membrane compacted structure crucial to ensure saltatory conduction and long term maintenance of axonal health. Thus, loss or damage of myelin is one of the major mechanisms underlying the neurodegenerative pathology observed in neurological disorders of diverse etiology, such as multiple sclerosis and leukodystrophies (Fancy *et al.*, 2011; Franklin and Ffrench-Constant, 2017; Saab and Nave, 2017). Spontaneous remyelination carried out by adult OPCs (aOPCs) can occur in demyelinated lesions and the newly formed myelin is able to restore at least partial physiological activity (Fancy *et al.*, 2011; Franklin and Ffrench-Constant, 2017). However, remyelination differs from the spatial and temporal organized program characteristic of developmental myelination (Fancy *et al.*, 2011), and occurs in the presence of an active immune response (Simons *et al.*, 2014; Miron, 2017). Although aOPCs differentiate into mature oligodendrocytes during remyelination the newly formed myelin is morphologically thinner for all but the smallest diameter myelinated axons (Fancy *et al.*, 2011; Franklin and Ffrench-Constant, 2017). Its capacity to prevent axonal degeneration in the long term has not been fully clarified (Saab and Nave, 2017). The molecular aspects associated with hypomyelination as a result of remyelination remain poorly understood, both in animal models of demyelination and in human patients (Fancy *et al.*, 2011; Franklin and Ffrench-Constant, 2017). In particular, whether and to what extent: 1) the remyelination proteome recapitulates that of developmental myelination, and 2) altered expression patterns of proteins within the newly formed myelin contribute to ineffective remyelination, are questions which remain to be addressed. Several studies have explored changes in the proteome of demyelinated lesions in a range of models, but without discerning on the cellular origin of the observed changes (Jahn *et al.*, 2009; Linker *et al.*, 2009; Werner *et al.*, 2010). Due to the complexity of the lesion environment, containing remyelinating glial cells, but strongly enriched in astrocytes and immune cells, no information on proteins which expression substantially changes in the myelin following remyelination is currently available. This is not trivial, as this knowledge could help identifying novel therapeutic strategies for a more efficient form of remyelination. Therefore, we addressed the above questions by specifically comparing the proteome of myelin formed after experimental demyelination (hereafter called the remyelination proteome) induced by injection of the toxin lysolecithin in the mouse spinal cord versus the proteome of myelin formed during development. Our data show that the remyelination proteome fails to recapitulate the one from developmental myelination presenting significant changes in the expression of modulators of metabolism, cell signaling and cytoskeleton remodeling proteins. We further show that downregulation of a single selected target is sufficient to partially hamper oligodendrocytes *in-vitro*.

## Materials and Methods

For list of antibodies, electron microscopy, iTRAQ mass-spectrometry, qRT-PCR, immunoblotting, immunohistochemistry, and quantification methods protocols see Supplementary Materials and Methods.

### Animals and demyelinating lesion model

All procedures were conducted with the approval and in accordance with the IBMC/i3S Animal Ethics Committee, the Portuguese Veterinary Office, the EU animal welfare laws, guidelines and policies, and following the ARRIVE guidelines. Wild type C57Bl/6 mice and Wistar rats of both sexes were used. Demyelination was obtained by focal injection of 1 μl 1% lysolecithin in the ventro-lateral funiculus of the spinal cord at vertebrae level T4. See Supplementary Materials and methods.

### Oligodendrocyte progenitor cell isolation and oligodendrocyte cultures

Oligodendrocyte progenitor cells were isolated from mix-glial cultures of P0-P2 Wistar Han rat brains, as described (Chen *et al.*, 2007). See Supplementary Materials and Methods.

### Viral-mediated shRNA

See Supplementary Materials and Methods.

### Myelinating co-cultures

See Supplementary Materials and Methods.

### Experimental design

Littermate and aged matched mice were randomly assigned to groups. No statistical method was used to predetermine sample size, but sample sizes are similar to those generally employed in the field. All quantifications were done blindly by third party concealment of treatments.

### Statistical analysis

All experiments were quantified blindly to the treatment. Statistics were analyzed using GraphPad Prism vs6.01. Data were assumed to be normally distributed, but not formally tested. Variance was assumed to be equal between groups. Statistical significance was determined using an unpaired two sample Student’s t-test for two group comparisons, while multiple group analysis was performed with one- or two-way analysis of variance (ANOVA) and post-hoc test as detailed in text/figures. Data show mean ± SEM of 3 independently run experiments unless otherwise specified. Significance was set at p < 0.05 *, p < 0.01 **, p < 0.001 ***.

### Data availability

The respective raw files used for iTRAQ data analysis were uploaded to a community data repository. Access credentials: ftp://MSV000085385@massive.ucsd.edu, MSV000085385, PWD: wlab@2020, Experiment 1: [ftp://MSV000085385@massive.ucsd.edu/raw/bdamaris_M1102_002.RAW], Experiment 2: [ftp://MSV000085385@massive.ucsd.edu/raw/bdamaris_M1102_004.RAW] Other data are available upon request from the corresponding author.

## Results

### The remyelination proteome differs from the proteome of developmentally formed myelin

We first addressed whether remyelination carries a distinct proteome by comparing proteins expressed in remyelination- versus developmentally-formed myelin (Fig. 1A, B). We induced focal demyelination in the thoracic spinal cord of adult C57Bl/6 mice by injection of lysolecithin (Fig. 1B, C and D). This model allows a temporal and spatial defined identification of a de-/re-myelinated lesion. Remyelination was largely complete at 21 dpi (Fig. 1B, C and D). Thus, we isolated myelin by sucrose gradient centrifugation using 1-cm of spinal cord tissue dissected around the lesion site at 21 dpi for the analysis of the remyelination proteome and from the intact myelinated spinal cord of young-adult mice for the analysis of the developmental myelin proteome (Fig. 2A). It has to be expected that a fraction of this tissue still contains normal myelin, surrounding the lesion area. Electron microscopy analysis confirmed specific enrichment in myelin membranes (Supplementary Fig. 1A). Comparison by SDS-PAGE and silver staining of total protein lysates obtained from myelin membranes isolated from remyelinated versus developmentally myelinated spinal cord tissue revealed the presence of numerous different bands (Supplementary Fig. 2A, arrows). To map the differentially regulated proteins, 2 independent biological samples per condition (consisting of spinal cord tissue pooled from 6-8 mice) were obtained and protein lysates labelled with iTRAQ reagents, followed by mass spectrometry and bioinformatic analysis (Fig. 2A, Supplementary Fig. 2B, C, and D). Applying an arbitrary 1.2-fold change threshold, we identified 103 proteins differentially expressed in myelin formed during remyelination when compared to a same total amount of myelin formed during development (Table 1, Supplementary Table 1). In setting this fold threshold, we took into consideration that differences in protein expression could be underestimated, due to remaining normal myelin in the dissected lesion-containing tissue. The great majority (97.1%) of regulated proteins showed decreased expression in the remyelination proteome compared to the naïve myelin proteome (Table 1, Supplementary Table 1). Only exceptions seem to be non-myelin proteins, probably derived from traces of contaminating blood (Hemoglobin-a and -b, Apolipoprotein A-II). Our results present relative changes, thus a strong fold change increase in a protein which is present in traces may have a far less biological significance than a small fold change decrease in a protein which is present at a much higher absolute quantity in the purified myelin samples (e.g. myelin specific proteins). We clustered the regulated proteins into functional categories by automated Panther analysis (www.pantherdb.org) (Fig. 2B, Table 1). Metabolism-related, cytoskeletal and signaling proteins were the main affected functional categories (Fig. 2B, Table 1). The remyelination proteome further revealed a myelin response (Fig. 2B, Table 1), characterized by lower expression of a subset of myelin specific proteins: MOG, MOBP, CNPase, and MAG. On the contrary, MBP showed comparable abundance in the remyelination versus the naïve myelin proteome, as also confirmed by immunoblotting (Supplementary Fig. 3 A and 4A). We also found dysregulation of proteins involved in fatty acid and cholesterol metabolism (Fig. 2B, Table 1). Literature mining revealed that some of these dysregulated targets were previously identified in the EAE mouse model and/or in plaques of multiple sclerosis patients. These included: Stathmin, Creatine-kinaseB, Sirtuin2, CRMP2 and 3 (Table 1, Supplementary Table 1) (Liu *et al.*, 2005; Jastorff *et al.*, 2009).

**Figure 1.**
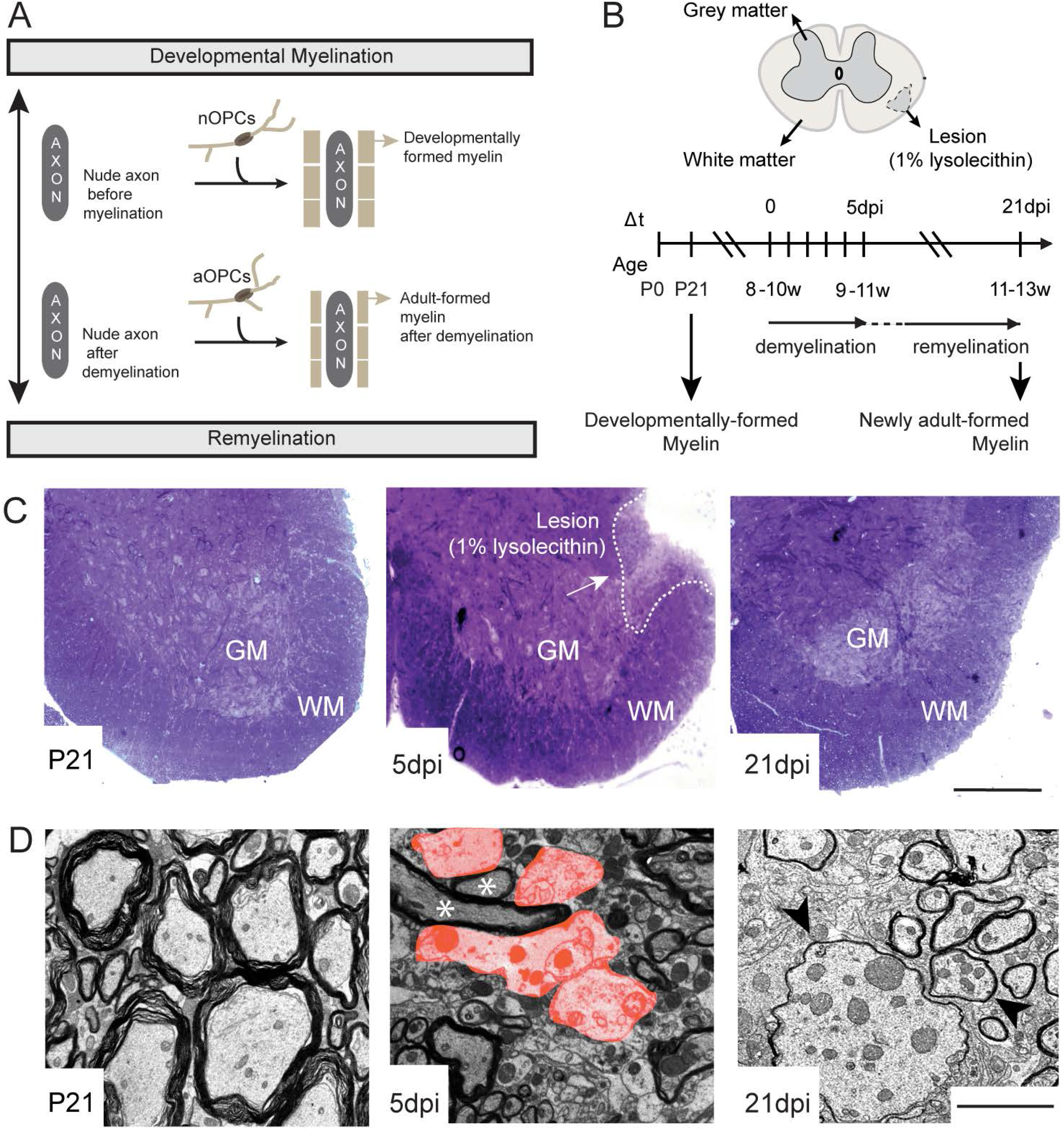
Experimental design. (**A**) Diagram illustrating the experimental design for the mapping of the changes in the proteome of myelin from the spinal cord following experimental de- and re-myelination. The proteome of myelin formed by aOPCs-derived oligodendrocytes, encasing axons following lysolecithin-induced focal demyelination, is compared to that of myelin formed by nOPCs-derived oligodendrocytes during development. (**B**) Timeline of lysolecithin-induced focal demyelination in the thoracic spinal cord (vertebrae level T4) and subsequent remyelination. Analyzed time-points are shown. (**C**) Representative toluidine blue staining of the ventral thoracic spinal cord at vertebrae level T4: postnatal day (P) 21 with axons invested by myelin, 5 days post-lysolecithin-injection (dpi) with a ventrolateral focal demyelinated lesion, and 21 dpi with axons reinvested by thinner myelin following remyelination. *n* = 3 mice observed per each time point. Scale bar: 250 μm (**D**) Representative micrographs depicting the morphology of the ventrolateral white matter by electron transmission microscopy: P21 with axons invested by myelin, 5 dpi with demyelinated axons in the lesion area (examples false colored in orange, examples of axons spared from demyelination indicated by asterisks), and 21 dpi with axons encased by thinner myelin following remyelination (examples indicated by arrow heads). *n* = 3 mice observed per each time point. Scale bar: 2.5 μm. OPCs = oligodendrocyte progenitor cells, dpi = days post injection, P = postnatal day, w = weeks, GM = grey matter, WM = white matter.

**Figure 2.**
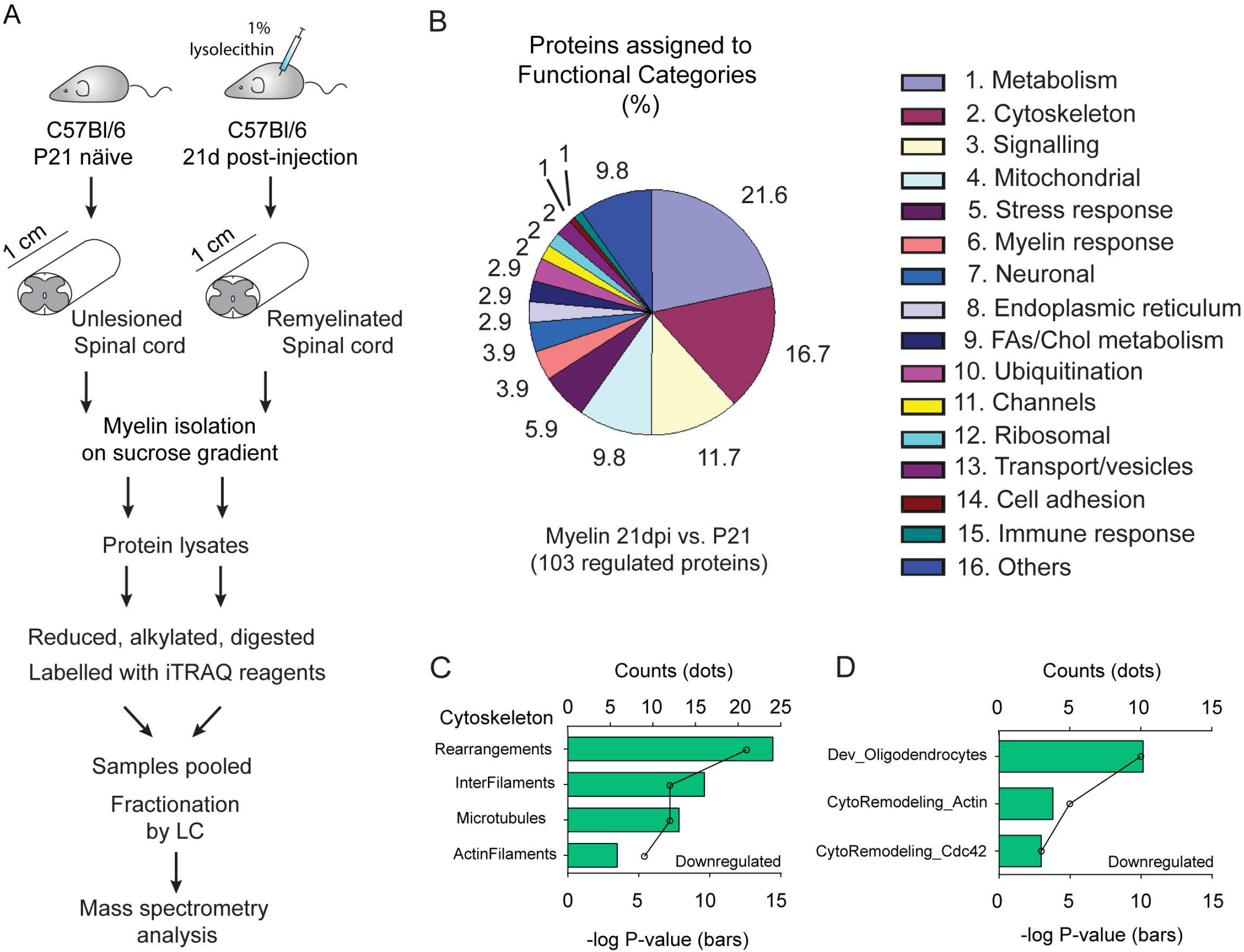
Analysis of the remyelination proteome reveals dysregulation of the cytoskeletal machinery. **(A)** iTRAQ experimental pipeline. 1 cm of spinal cord tissue around the lesion site (vertebrae level T4) was dissected from P21 naïve and 21 dpi C57Bl/6 mice for subsequent isolation of myelin on a sucrose gradient. Protein extracts were further processed for iTRAQ labelling and subsequent mass-spectrometry identification and relative quantification. **(B)** Diagram depicting percentage of dysregulated proteins based on assigned functional categories. *n* = 2 independent experiments, each with an independent set of pooled spinal cords for both 21 dpi and P21, each set from 6-8 mice (1cm of tissue around the lesion site or 1 cm from intact spinal cord tissue at the same thoracic spinal cord level). **(C, D)** Cytoskeletal-related Process Networks **(C)** and Pathway Maps (**D)** dysregulated in the remyelination proteome compared to the naïve myelin proteome, as identified by Metacore (version 6.29) enrichment analysis based on those proteins significantly downregulated in both independent experiments. Data points represent the total count of regulated proteins per each category (one value per category as output). Bar graphs represents the total −log P value per each category (one value per category as output).

**Table I.**
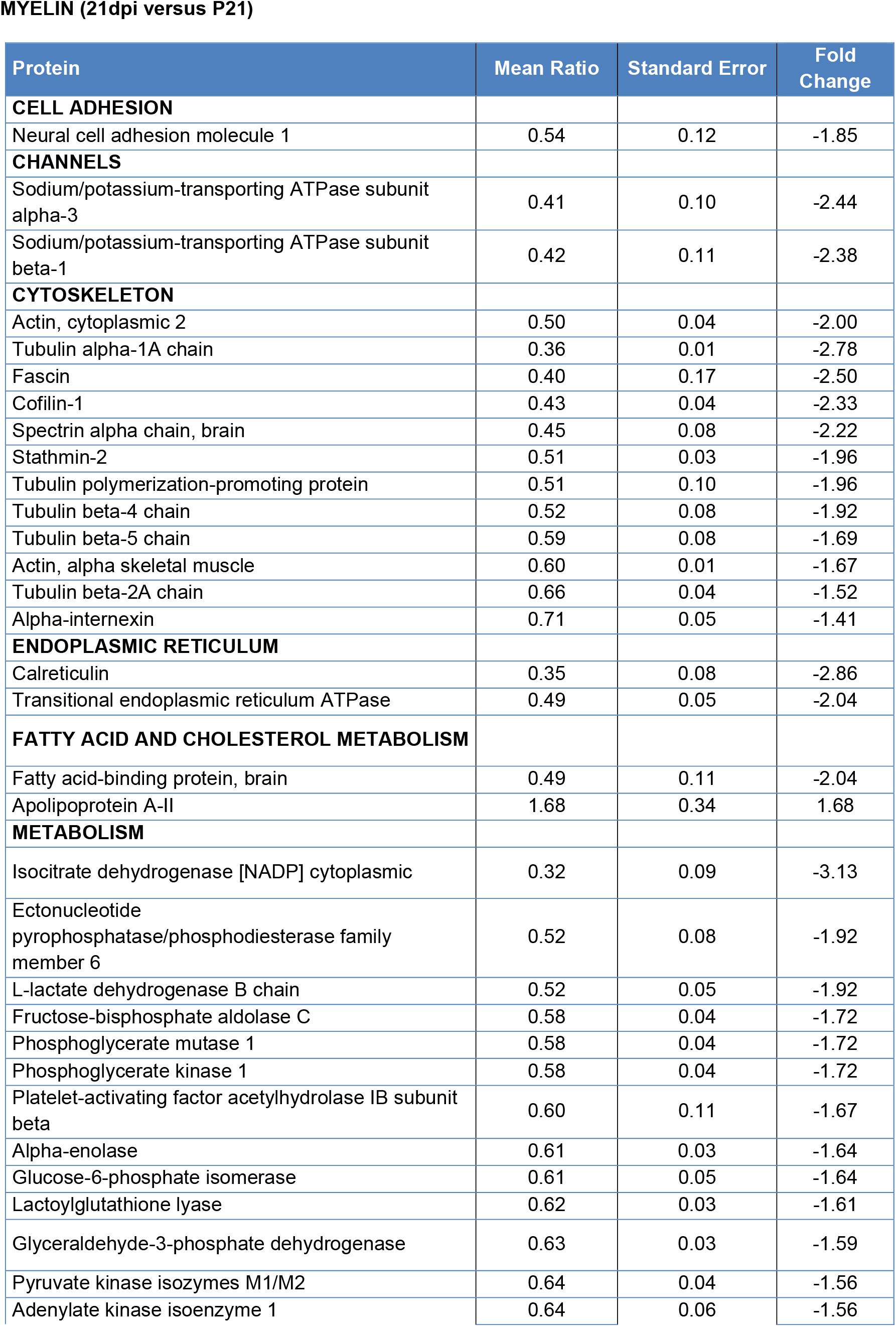
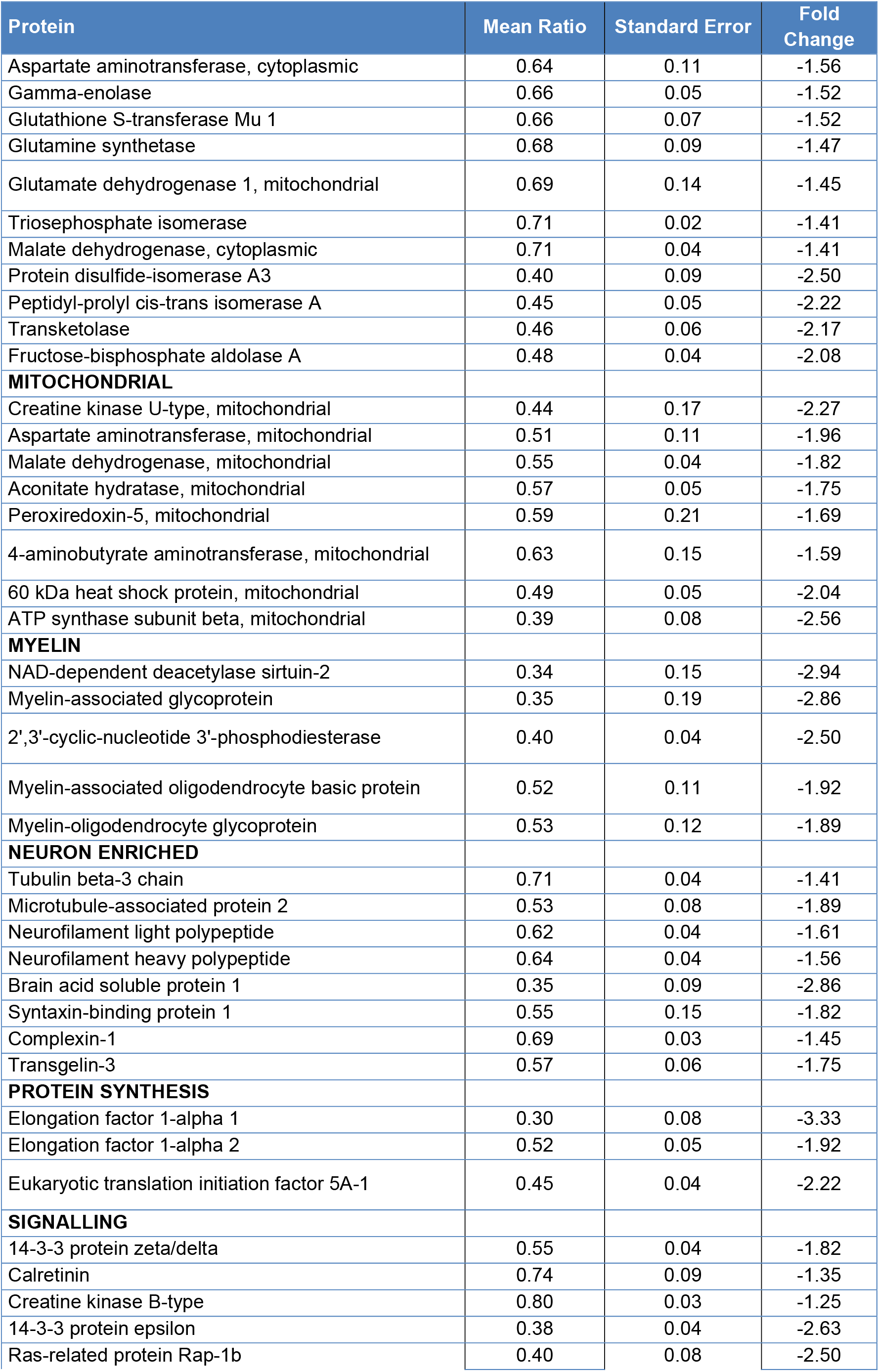
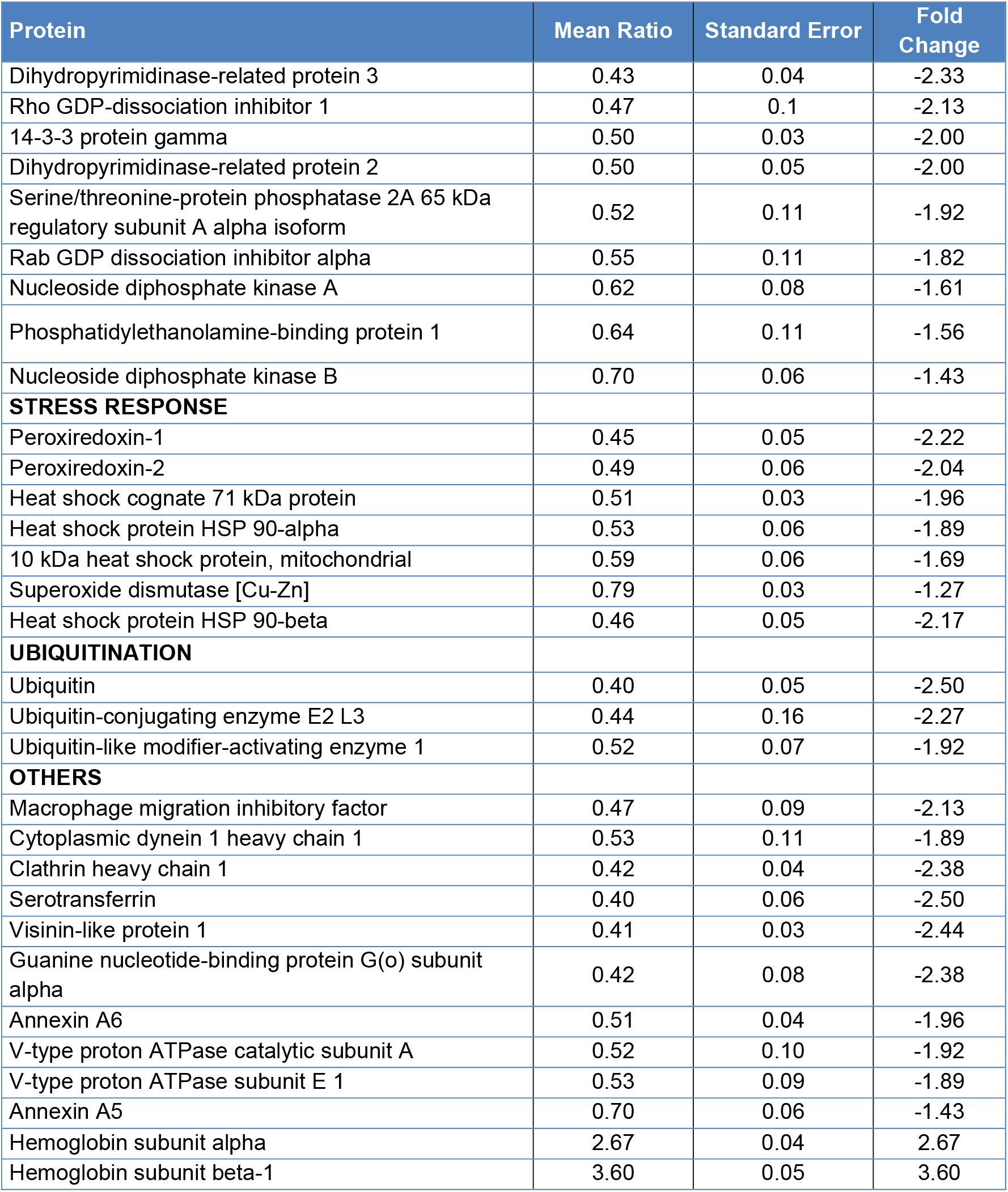
Proteins deregulated following remyelination, as identified by iTRAQ labelling followed by LC-MS/MS analysis.

**MYELIN (21dpi versus P21)**
| Protein | Mean Ratio | Standard Error | Fold Change |
| --- | --- | --- | --- |
| **CELL ADHESION** |  |  |  |
| Neural cell adhesion molecule 1 | 0.54 | 0.12 | -1.85 |
| **CHANNELS** |  |  |  |
| Sodium/potassium-transporting ATPase subunit alpha-3 | 0.41 | 0.10 | -2.44 |
| Sodium/potassium-transporting ATPase subunit beta-1 | 0.42 | 0.11 | -2.38 |
| **CYTOSKELETON** |  |  |  |
| Actin, cytoplasmic 2 | 0.50 | 0.04 | -2.00 |
| Tubulin alpha-1A chain | 0.36 | 0.01 | -2.78 |
| Fascin | 0.40 | 0.17 | -2.50 |
| Cofilin-1 | 0.43 | 0.04 | -2.33 |
| Spectrin alpha chain, brain | 0.45 | 0.08 | -2.22 |
| Stathmin-2 | 0.51 | 0.03 | -1.96 |
| Tubulin polymerization-promoting protein | 0.51 | 0.10 | -1.96 |
| Tubulin beta-4 chain | 0.52 | 0.08 | -1.92 |
| Tubulin beta-5 chain | 0.59 | 0.08 | -1.69 |
| Actin, alpha skeletal muscle | 0.60 | 0.01 | -1.67 |
| Tubulin beta-2A chain | 0.66 | 0.04 | -1.52 |
| Alpha-internexin | 0.71 | 0.05 | -1.41 |
| **ENDOPLASMIC RETICULUM** |  |  |  |
| Calreticulin | 0.35 | 0.08 | -2.86 |
| Transitional endoplasmic reticulum ATPase | 0.49 | 0.05 | -2.04 |
| **FATTY ACID AND CHOLESTEROL METABOLISM** |  |  |  |
| Fatty acid-binding protein, brain | 0.49 | 0.11 | -2.04 |
| Apolipoprotein A-II | 1.68 | 0.34 | 1.68 |
| **METABOLISM** |  |  |  |
| Isocitrate dehydrogenase [NADP] cytoplasmic | 0.32 | 0.09 | -3.13 |
| Ectonucleotide pyrophosphatase/phosphodiesterase family member 6 | 0.52 | 0.08 | -1.92 |
| L-lactate dehydrogenase B chain | 0.52 | 0.05 | -1.92 |
| Fructose-bisphosphate aldolase C | 0.58 | 0.04 | -1.72 |
| Phosphoglycerate mutase 1 | 0.58 | 0.04 | -1.72 |
| Phosphoglycerate kinase 1 | 0.58 | 0.04 | -1.72 |
| Platelet-activating factor acetylhydrolase IB subunit beta | 0.60 | 0.11 | -1.67 |
| Alpha-enolase | 0.61 | 0.03 | -1.64 |
| Glucose-6-phosphate isomerase | 0.61 | 0.05 | -1.64 |
| Lactoylglutathione lyase | 0.62 | 0.03 | -1.61 |
| Glyceraldehyde-3-phosphate dehydrogenase | 0.63 | 0.03 | -1.59 |
| Pyruvate kinase isozymes M1/M2 | 0.64 | 0.04 | -1.56 |
| Adenylate kinase isoenzyme 1 | 0.64 | 0.06 | -1.56 |
| Aspartate aminotransferase, cytoplasmic | 0.64 | 0.11 | -1.56 |
| Gamma-enolase | 0.66 | 0.05 | -1.52 |
| Glutathione S-transferase Mu 1 | 0.66 | 0.07 | -1.52 |
| Glutamine synthetase | 0.68 | 0.09 | -1.47 |
| Glutamate dehydrogenase 1, mitochondrial | 0.69 | 0.14 | -1.45 |
| Triosephosphate isomerase | 0.71 | 0.02 | -1.41 |
| Malate dehydrogenase, cytoplasmic | 0.71 | 0.04 | -1.41 |
| Protein disulfide-isomerase A3 | 0.40 | 0.09 | -2.50 |
| Peptidyl-prolyl cis-trans isomerase A | 0.45 | 0.05 | -2.22 |
| Transketolase | 0.46 | 0.06 | -2.17 |
| Fructose-bisphosphate aldolase A | 0.48 | 0.04 | -2.08 |
| **MITOCHONDRIAL** |  |  |  |
| Creatine kinase U-type, mitochondrial | 0.44 | 0.17 | -2.27 |
| Aspartate aminotransferase, mitochondrial | 0.51 | 0.11 | -1.96 |
| Malate dehydrogenase, mitochondrial | 0.55 | 0.04 | -1.82 |
| Aconitate hydratase, mitochondrial | 0.57 | 0.05 | -1.75 |
| Peroxiredoxin-5, mitochondrial | 0.59 | 0.21 | -1.69 |
| 4-aminobutyrate aminotransferase, mitochondrial | 0.63 | 0.15 | -1.59 |
| 60 kDa heat shock protein, mitochondrial | 0.49 | 0.05 | -2.04 |
| ATP synthase subunit beta, mitochondrial | 0.39 | 0.08 | -2.56 |
| **MYELIN** |  |  |  |
| NAD-dependent deacetylase sirtuin-2 | 0.34 | 0.15 | -2.94 |
| Myelin-associated glycoprotein | 0.35 | 0.19 | -2.86 |
| 2',3'-cyclic-nucleotide 3'-phosphodiesterase | 0.40 | 0.04 | -2.50 |
| Myelin-associated oligodendrocyte basic protein | 0.52 | 0.11 | -1.92 |
| Myelin-oligodendrocyte glycoprotein | 0.53 | 0.12 | -1.89 |
| **NEURON ENRICHED** |  |  |  |
| Tubulin beta-3 chain | 0.71 | 0.04 | -1.41 |
| Microtubule-associated protein 2 | 0.53 | 0.08 | -1.89 |
| Neurofilament light polypeptide | 0.62 | 0.04 | -1.61 |
| Neurofilament heavy polypeptide | 0.64 | 0.04 | -1.56 |
| Brain acid soluble protein 1 | 0.35 | 0.09 | -2.86 |
| Syntaxin-binding protein 1 | 0.55 | 0.15 | -1.82 |
| Complexin-1 | 0.69 | 0.03 | -1.45 |
| Transgelin-3 | 0.57 | 0.06 | -1.75 |
| **PROTEIN SYNTHESIS** |  |  |  |
| Elongation factor 1-alpha 1 | 0.30 | 0.08 | -3.33 |
| Elongation factor 1-alpha 2 | 0.52 | 0.05 | -1.92 |
| Eukaryotic translation initiation factor 5A-1 | 0.45 | 0.04 | -2.22 |
| **SIGNALLING** |  |  |  |
| 14-3-3 protein zeta/delta | 0.55 | 0.04 | -1.82 |
| Calretinin | 0.74 | 0.09 | -1.35 |
| Creatine kinase B-type | 0.80 | 0.03 | -1.25 |
| 14-3-3 protein epsilon | 0.38 | 0.04 | -2.63 |
| Ras-related protein Rap-1b | 0.40 | 0.08 | -2.50 |
| Dihydropyrimidinase-related protein 3 | 0.43 | 0.04 | -2.33 |
| Rho GDP-dissociation inhibitor 1 | 0.47 | 0.1 | -2.13 |
| 14-3-3 protein gamma | 0.50 | 0.03 | -2.00 |
| Dihydropyrimidinase-related protein 2 | 0.50 | 0.05 | -2.00 |
| Serine/threonine-protein phosphatase 2A 65 kDa regulatory subunit A alpha isoform | 0.52 | 0.11 | -1.92 |
| Rab GDP dissociation inhibitor alpha | 0.55 | 0.11 | -1.82 |
| Nucleoside diphosphate kinase A | 0.62 | 0.08 | -1.61 |
| Phosphatidylethanolamine-binding protein 1 | 0.64 | 0.11 | -1.56 |
| Nucleoside diphosphate kinase B | 0.70 | 0.06 | -1.43 |
| **STRESS RESPONSE** |  |  |  |
| Peroxiredoxin-1 | 0.45 | 0.05 | -2.22 |
| Peroxiredoxin-2 | 0.49 | 0.06 | -2.04 |
| Heat shock cognate 71 kDa protein | 0.51 | 0.03 | -1.96 |
| Heat shock protein HSP 90-alpha | 0.53 | 0.06 | -1.89 |
| 10 kDa heat shock protein, mitochondrial | 0.59 | 0.06 | -1.69 |
| Superoxide dismutase [Cu-Zn] | 0.79 | 0.03 | -1.27 |
| Heat shock protein HSP 90-beta | 0.46 | 0.05 | -2.17 |
| **UBIQUITINATION** |  |  |  |
| Ubiquitin | 0.40 | 0.05 | -2.50 |
| Ubiquitin-conjugating enzyme E2 L3 | 0.44 | 0.16 | -2.27 |
| Ubiquitin-like modifier-activating enzyme 1 | 0.52 | 0.07 | -1.92 |
| **OTHERS** |  |  |  |
| Macrophage migration inhibitory factor | 0.47 | 0.09 | -2.13 |
| Cytoplasmic dynein 1 heavy chain 1 | 0.53 | 0.11 | -1.89 |
| Clathrin heavy chain 1 | 0.42 | 0.04 | -2.38 |
| Serotransferrin | 0.40 | 0.06 | -2.50 |
| Visinin-like protein 1 | 0.41 | 0.03 | -2.44 |
| Guanine nucleotide-binding protein G(o) subunit alpha | 0.42 | 0.08 | -2.38 |
| Annexin A6 | 0.51 | 0.04 | -1.96 |
| V-type proton ATPase catalytic subunit A | 0.52 | 0.10 | -1.92 |
| V-type proton ATPase subunit E 1 | 0.53 | 0.09 | -1.89 |
| Annexin A5 | 0.70 | 0.06 | -1.43 |
| Hemoglobin subunit alpha | 2.67 | 0.04 | 2.67 |
| Hemoglobin subunit beta-1 | 3.60 | 0.05 | 3.60 |

In summary our proteomic analysis revealed that CNS remyelination leads to a distinct myelin protein composition, which does not fully recapitulate the one present in naïve developmentally-formed myelin.

### Altered expression of cytoskeleton remodeling proteins in the remyelination proteome

We next assessed whether observed differences in expression of proteins in the remyelination compared to the developmental myelin proteome could reflect a failure in molecular hubs critical for efficient myelination. We ran an unbiased Metacore (https://portal.genego.com/) enrichment analysis, which revealed inhibition of pathways and processes involved in the development of oligodendrocytes and in cytoskeleton remodeling (Fig. 2C, D). Modulation of actin- and tubulin-binding proteins downstream of Rho-GTPase signaling is critical towards efficient developmental myelination (Feltri *et al.*, 2008). In particular, tight regulation of actin dynamics is crucial in driving myelin growth (Nawaz *et al.*, 2015; Zuchero *et al.*, 2015). This led us to focus onto this proteins cluster.

Our analysis found the expression of RhoGDI, a key regulator of Rho-GTPase signaling, and of two of the main modulators of actin dynamics, namely CFL1/cofilin1 and FSCN1/Fascin1 to be downregulated in the remyelination proteome (Table 1 and Supplementary Table 1). Downregulation of FSCN1 (Fig. 3A and Supplementary 4C) and RhoGDI (Supplementary Fig. 3B and 4B) was further confirmed by immunoblotting. The phosphorylation levels of Ser-3, which regulate the function of CFL1 on actin were also diminished (Supplementary Fig. 3B and 4B). Our data show that these modulators of actin cytoskeletal dynamics in the remyelination proteome were significantly downregulated compared to the naïve myelin proteome.

**Figure 3.**
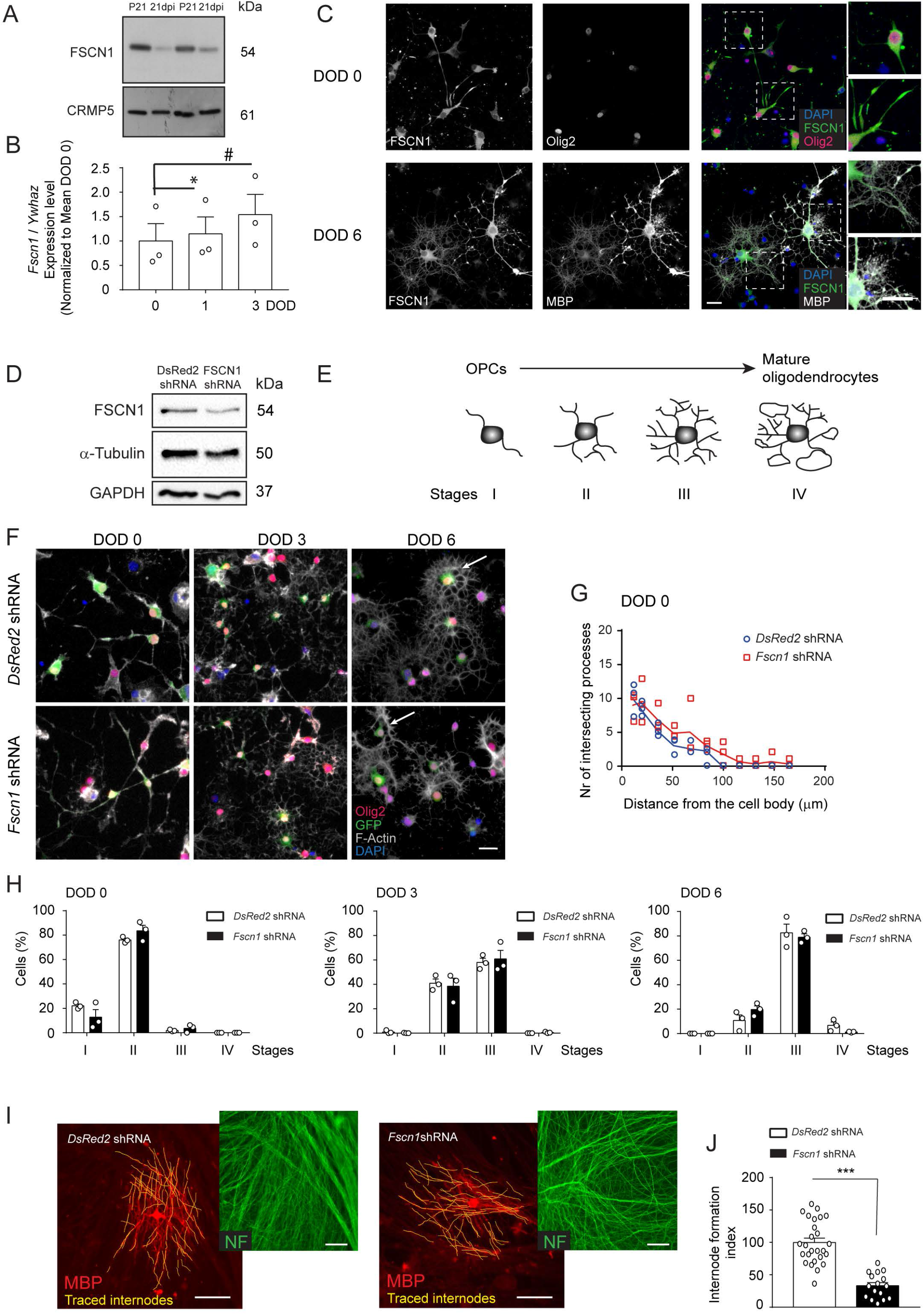
Consequences of decreasing FSCN1 expression in oligodendrocytes *in-vitro*. **(A)** Lower FSCN1 expression in newly adult-formed myelin following demyelination (21 dpi) compared to developmentally formed myelin (P21), shown by immunoblotting. Same amount of total proteins loaded (10 μg per lane). Due to their dysregulation at 21 dpi compared to P21, commonly used housekeeper proteins (GAPDH, Actin and Tubulin, see Table 1 and Supplementary Table 1) could not be used as loading control. CRMP-5 was not identified as regulated in the proteomic screening, and it is shown as reference protein for loading control as not being detected as regulated by immunoblotting. *n* = 2 independent sets of pooled spinal cords for both 21 dpi and P21, each from 8 mice (1cm of tissue around the lesion site or 1 cm from intact spinal cord tissue at the same thoracic spinal cord level). **(B)** Graph of qRT-PCR analysis for *Fscn1* at day of differentiation (DOD) 0, 1 and 3 in OPC-differentiating cultures. Data normalized to *Ywhaz* and to the mean of DOD 0. Data points represent *n* = 3 experiments (1-way Anova with Tukey’s multiple comparisons test DOD 0 vs. DOD 1 *df* = 2, *q* = 9.489, DOD 0 vs DOD 3 *df* = 2, *q* = 7.586), * p < 0.05, # p = 0.0597. **(C)** Representative immunostaining of differentiating-OPC cultures at 0 and 6 DOD. FSCN1 expression was detected in all Olig2+ oligodendrocytes already at 0 DOD. Note the prominent FSCN1 expression in fully mature promyelinating (MBP+) oligodendrocytes at 6 DOD. *n* = 3 observed experiments. Scale bar: 20 μm. **(D)** Cropped FSCN1 immunoblot with α-tubulin and GAPDH detected in the same membrane as loading controls. Quantification is shown in Supplementary Fig. 3C. From protein lysates obtained from OPCs infected with *Fscn1* shRNA or *DsRed2* shRNA, GFP+ sorted 48 hours following infection. Representative of *n* = 3 independent experiments. Full-length immunoblot in Supplementary Fig. 4. **(E, F)** Scheme of stage of differentiation along the oligodendrocyte-lineage (**E**) and representative immunostaining images (**F**) of cultures of GFP+ sorted Olig2+ OPCs treated with *Fscn1* shRNA or control *DsRed2* shRNA at 0, 3 and 6 DOD. Scale bar: 50 μm. **(G)** Graphic representation of Sholl Analysis’ results depicting the number of processes intersecting a radius at increasing distances from the cell body of oligodendrocytes. Data points represent *n* = 3 independent experiments (2-way Anova with Sidak’s multiple comparisons test *Fscn1* shRNA vs. *DsRed2* shRNA, *p* < 0.0001, *F_10,40_* = 31.82). **(H)** Corresponding bar graphs with quantification of percentage of cells assigned to the four different stages of oligodendrocyte differentiation, from OPC infected with *Fscn1* shRNA or control *DsRed2* shRNA, GFP+ sorted and differentiated in culture for 0, 3 and 6 DOD. Data points represent *n* = 3 independent experiments (two-way Anova with Sidak’s multiple comparisons test, *Fscn1* shRNA vs. *DsRed2* shRNA, DOD 0 stage I *p* = 0.1398, stage II *p* = 0.3005, stage III *p* = 0.9872, stage IV *p* > 0.9999, DOD 3 stage I *p* = 0.9998, stage II *p* = 0.9856, stage III *p* = 0.9785. stage IV *p* > 0.9999, DOD 6 stage I *p* < 0.9999, stage II *p* = 0.3060, stage III *p* = 0.9282, stage IV *p* = 0.7410). **(I)** Representative immunostaining of myelinating oligodendrocytes (MBP+) transduced with *Fscn1* shRNA or *DsRed2* shRNA as control, grown upon DRG neurites (NF+). Note the higher number of internodes formed by control oligodendrocytes (internodes traced in yellow). *n* = 2 independent experiments. Scale bar: 50 μm. **(J)** Corresponding bar graph showing the Internode formation index (= number of internodes per oligodendrocyte, normalized to the mean of the control set to 100) (unpaired two-tailed two sample Student’s t-test, *Fscn1* shRNA vs. *DsRed2* shRNA *p* < 0.0001, *t* = 7.282), *** p < 0.001. 9 to 16 cells analyzed per condition and experiment, in 2 independent experiments. Bars represent mean ± SEM. DOD = days of differentiation, OPCs = oligodendrocyte progenitor cells.

### Downregulating a single identified target is sufficient to partially impair oligodendrocytes *in-vitro*

Our screening identified changed expression of a plethora of proteins in the remyelination proteome. It is possible that the incomplete myelin formation of remyelination (thinner sheaths) in the lesioned CNS is the result of the synergistic effects of these concomitant changes. As an example, we reasoned that the downregulation in actin cytoskeleton-remodeling proteins might alter the dynamic of actin filament assembly/disassembly necessary for correct oligodendrocyte differentiation and/or myelin formation. To test if and to which degree dysregulation of a single identified target may impact on oligodendrocytes, we selected to address *in-vitro* the effect of downregulating the actin modulator FSCN1, which is transcribed at high levels in oligodendrocytes (http://www.brainrnaseq.org/).

We cultured neonatal rat-derived OPCs and induced them to differentiate *in-vitro* into mature oligodendrocytes. Quantitative real-time PCR confirmed that *Fscn1* transcription increased during *in-vitro* differentiation (Fig. 3B), and immunocytochemistry analysis showed FSCN1 protein to be expressed in oligodendrocytes (Olig2+), also at their mature MBP+ stage (Fig. 3C). We next studied the consequences of knocking down FSCN1 expression *in-vitro* in differentiating oligodendrocytes. We selected a validated shRNA sequence targeting *Fscn1*, based on our confirmation of its ability to decrease its expression (Fig. 3D). To test for a potential function of FSCN1 in regulating the morphological differentiation of oligodendrocytes, transduced cultured oligodendrocytes were categorized in 4 different classes based on the complexity of the branching of their processes, as previously described (Baer *et al.*, 2009) (Fig. 3E). At 0 DOD *Fscn1* shRNA-transfected OPCs showed longer processes (Fig. 3F). This observation was confirmed by Sholl analysis, with process length up to > 150 μm in *Fscn1* shRNA-transduced cells (Fig. 3G) compared to ~ 100 μm in control *DsRed2* shRNA-transduced oligodendrocytes (Fig. 3G). Despite this early effect, FSCN1 knock down did not significantly affect oligodendrocyte differentiation (Fig. 3H). In fact, the percentage of cells assigned to four categories of morphological differentiation at 0, 3 and 6 DOD *in-vitro* in both control and *Fscn1* shRNA-transduced cells were not significantly different (Fig. 3H). Furthermore, at 6 DOD no significant differences were found between controls and FSCN1 knockdown OL concerning the percentage of mature (MBP+ lamellae-forming) oligodendrocytes (Supplementary Fig. 5A), or in their capacity of forming myelin-like sheaths (Supplementary Fig. 5B).

Next, to investigate whether reducing FSCN expression impacted on the capacity of oligodendrocytes to myelinate axons, we plated FACS-sorted GFP-expressing *Fscn1* shRNA-transduced oligodendrocytes (48 hours post infection) onto purified naïve dorsal root ganglia neurons. After 18 days in culture, oligodendrocytes transduced with control shRNA had myelinated axons forming several myelin internodes (Fig. 3I). In contrast, oligodendrocytes in which FSCN1 had been knocked down displayed a lower number of internodes per cell (Fig. 3I and 3J), with a slightly increased length (Supplementary Fig. 5C).

In summary, our data show that knocking down a single target – altered in the remyelination proteome compared to the myelination proteome – may be sufficient to impact upon oligodendrocyte normal characteristics.

## Discussion

In this study, we assessed whether the remyelination proteome differs from the developmental myelin proteome. Remyelinating tissue contains high density of astrocytes, microglia and lymphocytes that proliferate and get activated at the lesion site. Thus, using as input myelin contaminated with membranes from these cells is likely to mostly identify the upregulation of markers for gliosis in a proteomics comparison of lesioned versus intact tissue. A contamination with astrocytes, microglia and lymphocytes and their respective proteomes was prevented by applying a myelin-isolation protocol. No marker for gliosis was found to be upregulated. Our proteomic analysis of myelin isolated from lesioned versus intact tissue showed that the proteome of myelin formed following remyelination is distinct from that of the myelin formed during development. Based on these results and functional *in-vitro* assays, we propose that changes in the expression of key modulators of actin cytoskeleton dynamics might contribute to restrict the extent of remyelination.

Several lines of evidence established that regulation of actin dynamics is key in driving oligodendrocyte differentiation and myelination during development (Nawaz *et al.*, 2015; Samanta and Salzer, 2015; Zuchero *et al.*, 2015; Shao *et al.*, 2017; Azevedo *et al.*, 2018). Our data provide evidence that actin binding proteins are dysregulated in myelin formed following de-/re-myelination and that inefficient expression of a single actin bundling protein, FSCN1, is sufficient to at least partially impact upon oligodendrocytes normal morphology *in-vitro*. This shows that our approach is suitable for a first identification of novel potential therapeutic targets, which will require further validation *in-vivo* in future studies. Nonetheless, the changes observed upon *in-vitro* knock down of a single target, i.e. FSCN1, were not dramatic. This lead us to conceive that inefficient remyelination is likely to be driven by the concomitant dysregulation of several proteins. Identifying the molecular mechanisms leading to their lower expression in remyelinating oligodendrocytes remains a goal for future studies.

We were surprised to find downregulation of major myelin proteins in the remyelination proteome compared to the naïve myelin proteome. We cannot rule out that the presence of aOPC-derived Schwann cell remyelination (Zawadzka *et al.*, 2010) may partially account for the observed lower expression of CNS myelin-specific proteins, MOG and MOBP. However, this could not explain the decrease in MAG and CNPase, also expressed by Schwann cells. MBP, another major myelin protein expressed by both oligodendrocytes and Schwann cells, showed no significant difference in its total expression in the remyelination versus the myelin naïve proteome. Nonetheless immunoblotting revealed diverse level of expression of its different isoforms. Our data may indicate dysregulation of specific myelin proteins following remyelination in the applied lesion model. This finding was unexpected as these proteins are critical to maintain the highly compacted organization of myelin membranes (Bakhti *et al.*, 2014), thus highlighting the importance of future studies addressing the long term stability of remyelinated profiles. It was surprising that 97% of the differentially expressed proteins identified displayed lower expression (by more than 1.2 fold) in remyelinated membranes compared to myelinated membranes. It must be considered that the samples may be in different “developmental” stages, i.e. P21 myelin is fully mature/homeostatic whereas 21dpi myelin is still undergoing maturation. This could account for the observed diverse expression of myelin specific proteins. Applying a similar experimental approach to later time points, in future studies, will allow to address if and when myelin formed in the remyelination process is capable of reaching maturity.

In the course of our study we found that proteins involved in the metabolism of cholesterol and fatty acids showed a diverse expression in the remyelination versus the naïve myelin proteome. Myelination relies on the production of a massive amount of cell membranes which contain a high percentage of lipids. Accordingly, both fatty acid synthesis and cholesterol are critical for myelination during development (Saher *et al.*, 2005; Schmitt *et al.*, 2015; Camargo *et al.*, 2017; Montani *et al.*, 2018; Dimas *et al.*, 2019). It is therefore conceivable that an impairment in lipid synthesis and/or transport could contribute in restricting myelin growth during remyelination, an aspect that warrants further attention. We cannot currently exclude that control versus remyelinated samples may present a diverse quality of myelin enrichment in the myelin isolated fraction, e.g. due to a diverse solubility of myelin membranes from remyelinated tissue due to a different lipid composition. This is an interesting and complex aspect which remains to be further addressed.

## Abbreviations

dpi: days post injury
DOD: days of differentiation
iTRAQ: isobaric Tags for Relative and Absolute Quantitation
OPCs: Oligodendrocyte Progenitor Cells
aOPCs: adult OPCs
nOPCs: neonatal OPCs
P(n): Postnatal day (n)
PNS: peripheral nervous system

## Acknowledgments

We thank members of the Relvas lab for discussion and the Scientific Platforms of IBMC/i3S for excellent technical support.

## Supplementary Material

Supplementary material is available as separate .pdf file.

