## Supplemental Material for "The proteome of remyelination is different from that of developmental myelination"

### Supplementary Materials and Methods

#### *Antibodies*

Primary antibodies: rabbit anti-p(S3)-CFL1/cofilin1 (WB 1:500, Cat# ab12866, Abcam, Cambridge, UK), anti-CRMP5 (WB 1:2000, Cat# ab36203, Abcam, Cambridge, UK), mouse anti-FSCN1/fascin1 (WB 1:400, ICC 1:50, clone SPM133, Cat# ab15110, Abcam, Cambridge, UK), mouse anti-GAPDH (WB 1:15000, Cat# 5G4, HyTest, Turku, Finland), rabbit anti-Olig2 (ICC 1:500, Cat# AB9610, Merck Millipore, Burlington, MA, USA), rat anti-MBP (WB 1:1000, ICC 1:100, clone 12, Cat# MCA409S, Biorad, Hercules, CA, USA), rabbit anti-RhoGDI (WB 1:1000, Cat# 2564, Cell Signaling Technologies, Danvers, MA, USA), mouse anti-αTubulin (WB 1:40000, clone B-5-1-2, Cat# T5168, Sigma, St. Louis, MO), rabbit anti-βIIITubulin (ICC 1:1000, Cat# 302302,Synaptic Systems, Göttingen, Germany). Secondary antibodies: goat anti-mouse Alexa488 (ICC 1:1000, Cat# A11001, Molecular Probes, Thermo Fisher Scientific, Rockford, IL, USA), goat anti-rat Alexa568 (ICC 1:1000, Cat# A11077, Molecular Probes, Thermo Fisher Scientific, Rockford, IL, USA), goat anti-rabbit Alexa568 (ICC 1:1000, Cat# A11011, Molecular Probes, Thermo Fisher Scientific, Rockford, IL, USA), goat anti-mouse Alexa647 (ICC 1:1000, Cat# A21235, Molecular Probes, Thermo Fisher Scientific, Rockford, IL, USA), goat anti-rat Alexa647 (ICC 1:1000, Cat# A21247, Molecular Probes, Thermo Fisher Scientific, Rockford, IL, USA), goat anti-mouse Cy3 (ICC 1:1000, Cat# 115-165-146, Jackson Laboratory, Bar Harbor, ME, USA), goat anti-mouse HRP (WB 1:10000, Cat# 115-035-146, Jackson Laboratory, Bar Harbor, ME, USA), goat anti-rabbit HRP (WB 1:10000, Cat# 111-035-144, Jackson Laboratory, Bar Harbor, ME, USA). DAPI (4’,6’-diamidino-2-phenylindole dilactate) (Cat# D3571, Thermo Fisher Scientific, Rockford, IL, USA) was used for nuclear staining.

#### *Animals and demyelinating lesion model*

All procedures were conducted with the approval and in accordance with the IBMC/i3S Animal Ethics Committee, the Portuguese Veterinary Office, the European Union animal welfare laws, guidelines and policies, and following the ARRIVE guidelines. Wild type C57Bl/6 mice (*Mus musculus*) and Wistar rats (*Rattus norvegicus*) of both sexes were used. Animals were group-caged and kept in a 12 hours' light/dark cycle, at 22 ± 2 ºC, with ad libitum access to food and water. Littermates and aged-matched animals were randomly assigned to experimental groups. Demyelination by focal injection of 1 µl of 1% lysolecithin (Cat# L4129, Sigma, St. Louis, MO) in the ventro-lateral funiculus of the spinal cord at vertebrae level T4 was performed. Briefly, animals received a single pre-operative subcutaneous injection of Buprenorphine (0.05 mg/kg) as analgesic and were anaesthetized by intraperitoneal injection of Medetomidine (75 mg/kg) and Ketamine (1 mg/kg). A dorsal laminectomy was performed and the dura pierced with an acupuncture needle. 1 µl of 1% lysolecithin (vol/vol) in 1x PBS, pH 7.4, was injected hemilaterally in the ventro-lateral funiculus at a rate of ~ 0.5 µl per minute with a glass needle coupled to a Hamilton syringe, via a three-way micromanipulator (Narishige, Japan). The overlaying musculature and skin were sutured. Animals were injected with Atipamezole (1 mg/kg) to revert from anesthesia and let to recover on a heating pad at 37°C. Animals received a subcutaneous injection of 0.1 mg / kg of Buprenorphine as analgesic the two days following the injury.

#### *Electron microscopy*

Mice received an intraperitoneal terminal injection of pentobarbital (150mg/kg) and were intracardially perfused with 0.1M phosphate buffer (PB) pH 7.4 followed by EM fix solution (2.5% glutaraldehyde, 4% paraformaldehyde, in 0.1 M PB pH 7.4). Spinal cords were dissected and a 1 cm of spinal cord tissue around the lesion area or from intact spinal cord tissue at vertebral level T4 was isolated and postfixed in EM fix solution overnight at 4°C. After dehydration through an acetone series, tissues were post-fixed in 2% osmium tetroxide overnight, and embedded in Spurr's resin (Electron Microscopy Sciences, Hatfield, PA, USA). 65 nm-thick ultrathin sections were cut on a Reichert-Jung Ultra cut E ultramicrotome (Leica, Germany), transferred onto copper grids with a carbon film (Electron Microscopy Sciences, Hatfield, PA, USA), and counterstained with 2% uranyl acetate and 1% lead citrate. Sections were observed in a transmission electron microscope (Jeol 902A model) (Carl Zeiss Oberkochen, Germany) at 80 kV. Images were digitally recorded using a Gatan SC 1000 ORIUS CCD camera (Warrendale, PA, USA).

*Myelin isolation*

Two separate independent experiments were performed, each with independent biological samples. Briefly, 6 to 8 mice per group (i.e. treatment and experiment) were sacrificed and 1-cm of thoracic spinal cord tissue was dissected: (i) around the lesion site in mice which underwent focal demyelination or (ii) the corresponding level in the intact spinal cord of naïve mice. The 1-cm dissected tissue was measured as 0.5-cm on both sides from the point of injection of lysolecithin. This was identified upon dissection by the presence of a small scar on top of the injection area. Tissue fragments were peeled off of the meninges and contaminating blood, and they were pooled for condition (remyelinated tissue versus intact myelinated tissue) and further processed for myelin purification, as previously described (Larocca and Norton, 2007). Briefly, spinal cords were transferred in suspension buffer (0.32 M sucrose in 50 mM Tris-HCl pH 7.5, 5 mM EDTA) and tissue disrupted in a glass and stainless steel manual homogenizer. The homogenate was layered onto a 0.83 M sucrose suspension buffer, and centrifuged at 140000g in a SW41Ti rotor (Beckman, Fullerton, CA, USA) for an hour. The obtained white band of crude myelin membranes was collected and subjected to three cycles of osmotic shock in 50 mM Tris-HCl pH 8.3, 5mM EDTA, 1x at 75000 g and 2x at 12000 g at 4 ºC. Myelin pellets were resuspended and subjected to a repetition of density centrifugation and osmotic shock. Finally, obtained myelin pellets were resuspended in 0.83 M sucrose suspension buffer overlaid with 0.32M suspension buffer and centrifuged at 75000 g for 30 min at 4°C. The myelin fraction was subjected to a last repetition of osmotic shock and the pellet either processed for electron microscopy analysis according to the same protocol describe above for tissue or resuspended in 1% Rapigest, 50mM Tris-HCl pH 7.5, 2mM EDTA, for proteomic analysis.

#### *iTRAQ labeling*

For subsequent iTRAQ labeling, proteins were purified by acetone precipitation and resuspended in the iTRAQ kit (Applied Biosystems, Foster City, CA, USA) dissolution buffer supplemented with 0.2% Rapigest surfactant (Waters, Milford, MA, USA). Protein concentration was determined by Nanodrop. 150 µg proteins per each condition (P21 and 21dpi) and experiment were used for subsequent labelling according to the manufacturer's protocol. Briefly, proteins were denatured and reduced, cysteines were blocked and subsequently proteins digested overnight with trypsin (Promega, Madison, Wi, USA). Peptide samples were combined and labeled with iTRAQ reagents (reagent 115 for P21 samples and reagent 117 for 21dpi samples). Two other channels (reagent 114 and 116) were used in relation to another experiment which was not followed up upon in the context of this manuscript. Following, labeled peptides were purified by Cation-Exchange Chromatography and desalted through a Sep-Pak C18 cartridge according to manufacturer's protocols. Samples were dried in a SpeedVac and peptides solubilized in 100ul HPLC-grade water, 5% acetonitrile, 0.1% formic acid.

#### *Peptide Analysis by LC-MS/MS*

Peptide mixtures were analyzed on a LTQ-Orbitrap XL mass spectrometer (Thermo Fisher Scientific, Rockford, IL, USA). Peptides (approximately 500 ng/injection) were loaded on a 10 cm reversed phase-high performance liquid chromatography column (75 μm diameter) packed with C18 material (Magic C18 AQ 3 μm; Michrom Bioresources, Auburn, CA, USA) and eluted into the mass spectrometer over a linear gradient of 5–30% Buffer B (2% H2O, 0.1% formic acid in acetonitrile) in Buffer A (2% acetonitrile, 0.1% formic acid in H2O) for 60 min. The flow rate was set to 0.3 μL/min. For the analysis of iTRAQ-labeled peptides, the instrument was operated in parallel mode, allowing accurate mass measurement of the four most intense precursor ions (350-1600 m/z) in the ion precursors in the Orbitrap concurrent with the acquisition of data-dependent CID MS/MS. Precursors with a minimal count of 250 were first fragmented by CID MS/MS and then again fragmented with a HCD MS/MS scans (with 7500 FWHM resolution at m/z=400). Peptide precursor ions were selected with an isolation window of 2 Da, and a target value of 1×105. Dynamic exclusion was implemented with a repeat count of 1 and exclusion duration of 30 s. The NCE was set to 35% for CID, and 55% for HCD. Precursors with charge 1 were excluded from the analysis.

#### *Database search and iTRAQ quantification*

Raw data files were converted to the open mzXML data format (Pedrioli *et al.*, 2004) by the ReAdW algorithm (v4.3.1), and scans were renumbered ensuring the appearance of each scan number only once using an in-house script. MS peptide spectra were searched by the SEQUEST against the human UniProt-SwissProt protein database (version 57.15) extended by common protein contaminants and reversed sequences as decoy entries (in total 40,521 protein entries). Amino acid modifications were permitted for static carbamidomethylation of cysteine (+57.021 Da) and variable oxidation of methionine (+15.995 Da). Peptide mass tolerance was set to 0.04 Da. Search allowed for semitryptic peptides and one missed peptide cleavage. Statistical evaluation was based on a target-decoy search strategy and was performed by software tools derived from the Trans Proteomic Pipeline (TPP version 4.4) (Keller *et al.*, 2005). A peptide and protein false discovery rate of 1% was determined using the nonparametric model in combination with the decoy option in Peptide Prophet. Quantification of the iTRAQ signal and the calculation of ratios was performed with the software LIBRA, included in the TPP.

Data were exported to Excel to calculate differential expression of proteins in 21dpi versus P21 samples. Only proteins identified by ≥ two unique peptides and with an identification probability score ≥ 0.9 were considered significantly regulated. A 1.2-fold change threshold was applied for both up and down regulations. Only proteins which passed this threshold and were found significantly up and down regulated in both independent experiments were used for identification of candidates, clustering in functional categories and reported in Fig. 2, Table 1 and Supplementary Table 1.

#### *Estimation of iTRAQ labeling efficiency*

The mzXML files from the two iTRAQ measurements were searched with Comet (version 2018.01 rev. 2) while allowing for a variable modification of 144.10253 Da at K residues and N-termini. XPRESS (TPP v5.2.0-rc5) was used to integrate the elution profiles of labeled and unlabeled precursor ions. The list of identified peptides was filtered at a 1% FDR (as calculated by PeptideProphet (TPP v5.2.0-rc5)) and finally the labeling efficiency was calculated using python (3.7.6) as

$$labelingPercentage = \frac{heavyPeptideArea}{lightPeptideArea + heavyPeptideArea}$$

#### *Silver staining*

Gels were fixed at room temperature for 1 hour in 20% ethanol, 5% acetic acid, then shortly bathed in Farmers' solution (0.05% Na2CO3 *3H2O, 0.15% K3CN6Fe, 0.30% Na-thiosulfate) and washed with $H_2O$. After incubation for 20min in 0.2% silver nitrate solution, staining was developed with 3% $Na_2CO_3$, 0.05% formaldehyde.

*Immunoblotting*

Protein lysates were prepared in 50mM Tris-HCl pH 8.3, 5mM EDTA, 0.1% RapiGest SF surfactant (Waters, Milford, MA, USA) for confirmation of proteomic data, or in RIPA lysis buffer (50 mM Tris pH 8.0, 150 mM NaCl, 1% NP-40, 1 mM EDTA, 0.5% sodium deoxycholate, 0.1% sodium dodecyl sulfate (SDS) supplemented with protease inhibitor cocktail (Sigma)) for confirmation of FSCN1 knockdown. Protein concentration was measured by Nanodrop and an equal amount of proteins loaded per lane and resolved onto NuPAGE (Invitrogen, Carlsbad, CA, USA) or SDS-PAGE gels. Resolved proteins were transferred on PVDF Immobilon-P (Millipore, Bedford, MA, USA) or Nitrocellulose (GE Healthcare, Life Sciences) membranes overnight. Membranes were stained with 0.1% Coomassie R-250 (Sigma, St Louis, MO, USA) in 40% methanol for 20 seconds or with Ponceau S staining solution (Sigma, St Louis, MO, USA) for 2-3 minutes to confirm equal protein loading between lanes. Correctly loaded membranes were de-stained, from Coomassie staining with 3x washing in 50% methanol, from Ponceau staining with 1x $H_2O$ washing and 1x 0.1M NaOH washing, and further processed. Membranes were blocked in 3% BSA, 0.1% Tween-20, 1X PBS, for detection of phosphorylated cofilin, 5% milk powder, 1X TBS for detection of FSCN1, or 3% milk powder, 0.1% Tween-20, 1X PBS for detection of other proteins. Incubation with primary antibody overnight at 4°C in blocking buffer and, subsequently, secondary HRP-conjugated antibody for 1 hour at room temperature, followed. For detection of CRMP-5, membranes used for detection of FSCN1 were treated with Restore PLUS Western Blot Stripping Buffer (Thermo Fisher Scientific) and re-probed. Signals were detected with SuperSignal WestPico chemiluminescent substrate (Pierce, Thermo Fisher Scientific,

Rockford, IL, USA). Chemiluminescent signal was: i) detected with RX-films (Fujifilm Europe, Düsseldorf, Germany) scanned on a Molecular Imager GS800 calibrated densitometer (Bio-Rad Laboratories, Hercules, CA, USA), and images acquired with Quantity-1D v.4.6 software (Bio-Rad Laboratories, Hercules, CA, USA), or ii) detected and imaged with a ChemiDoc XRS System (Bio-Rad Laboratories, Hercules, CA, USA). Only images containing bands with no saturated pixels, confirmed by the software detection systems, were used for subsequent quantification. Densitometry and quantification of the relative levels were carried out with ImageJ or Image lab software vs5.2.1 (Bio-Rad Laboratories, Hercules, CA, USA) software.

#### *Oligodendrocyte progenitor isolation and oligodendrocyte cultures*

Oligodendrocyte progenitor cells were isolated from mix-glial cultures of P0-P2 Wistar Han rat brains, as previously described (Chen *et al.*, 2007) . Briefly, cortices from 10-12 animals were dissected in ice-cold Hank's Balanced Salt Solution (HBSS) (Gibco, Thermo Fisher Scientific, Rockford, IL, USA) with 1% penicillin / streptomycin (Thermo Fisher Scientific, Rockford, IL, USA), meninges removed and tissue mechanically homogenized. The tissue suspension was digested in 0.25% Trypsin-EDTA (Gibco, Thermo Fisher Scientific, Rockford, IL, USA), 0.1 mg/mL DNase I (Sigma, St Louis, MO, USA) in HBSS for 15 minutes at 37 °C, and the cells pellet resuspended in 2 volumes of complete DMEM (c-DMEM) (Cat# 31966047, Invitrogen, Thermo Fisher Scientific, Rockfore, IL, USA) supplemented with 10% fetal bovine serum (FBS) (Gibco, Thermo Fisher Scientific, Rockford, IL, USA) and 1% penicillin/streptomycin (Thermo Fisher Scientific, Rockford, IL, USA). After centrifugation, cells were resuspended in c-DMEM and filtered through a 100 µm nylon cell strainer (Falcon, Corning, Tewksbury, MA, USA). Cells were plated on PDL (poly-D-lysine, Cat# P7405, Sigma, St Louis, MO, USA) coated T75 flasks (Sarstedt, Nümbrecht, Germany) at a density of ~2 brains per flask and incubated at 37°C, 5%

$CO_2$ for 10 days. Medium was replaced every 2-3 days. To enrich in oligodendrocyte progenitor cells, flasks were shaken at 220 rpm for 2 hours in a Minitron incubator (Infors HT, Annapolis Junction, MD, USA) at 37°C to remove microglial cells. After addition of fresh c-DMEM, the flasks were shaked at 240 rpm overnight at 37°C. The following day, the medium was collected into Petri dishes and incubated for 2 hours, to eliminate adhering cells. The remaining cell suspension was passed through a 40 µm nylon cell strainer (Falcon, Corning, Tewksbury, MA, USA) and plated onto acid-washed PDL coated glass coverslips (2.5 µg/ml PDL diluted in boric acid buffer [50mM boric acid, 12.5 mM sodium tetraborate) at a density of 45'000 cells per 14-mm of diameter coverslip or 3 x $10^5$ cells/well in 6-well plates in SATO medium (5 mg/ml insulin (Sigma, St Louis, MO, USA), 100 µg/ml human apo-transferrin (Sigma, St Louis, MO, USA), 100 µg/ml bovine serum albumin (NZY Tech, Lisbon, Portugal), 60 ng/ml progesterone (Sigma, St Louis, MO, USA), 16 µg/ml putrescine (Sigma, St Louis, MO, USA), 40 ng/ml sodium selenite (Sigma, St Louis, MO, USA), 40 ng/ml thyroxine (Sigma, St Louis, MO, USA), 30 ng/ml thriiodo-L-thyronine (Sigma, St Louis, MO, USA) in DMEM). SATO medium was initially supplemented with 10 ng/ml PDGFaa (Prepotech, London, UK) and 10 ng/ml FGF (Prepotech, London, UK). To induce oligodendrocyte differentiation, PDGFaa and FGF were withdrawn and 0.5 % FBS added.

*Viral-mediated shRNA*

The validated shRNA sequence #TRCN0000108929 (Sigma, St Louis, MO, USA) targeting mouse *Fscn1* was purchased as shRNA glycerol stock. Plasmids were produced according to manufacturer's protocol and purified with a Qiagen Plasmid Midi Kit. As control shRNA, a sequence targeting *dsRed2* (Ozcelik *et al.*, 2010) was used: 5'-AGTTCCAGTACGGCTCCAA-3'. The sequences were cloned into the pSicoR vector (Ozcelik *et al.*, 2010), which also carries the expression sequence for GFP (green fluorescent protein), using HpaI-XhoI cloning sites. Bacteria clones carrying the construct were sequence verified. The vector was then used for high titer lentivirus production. Briefly, HEK293T cells cultured on 10 cm PDL-coated dishes (~95 % confluent) were transfected using jetPrime reagent (Polyplus-transfection, New York, USA) with pSicoR vector containing the target sequence and the viral packaging constructs psPAX2 and VSVG, according to the manufacturer's instructions. 48 hour later the viral particles were purified and concentrated using Amicon Ultra 15 ml centrifugal filters.

Upon seeding, oligodendrocyte progenitor cells were transduced with lentiviral particles (multiplicity of infection ratio (MOI) ≥1). Infection efficiency was assessed by GFP expression.

*Myelinating co-cultures*

Myelinating co-cultures were prepared with minor modifications to previously described protocols (Azevedo *et al.*, 2018). Briefly, embryonic (E) dorsal root ganglia (DRG) neurons were isolated from E15 rats and dissociated in 0.05 % Trypsin-EDTA and 0.4 mg/ml DNase for 90 minutes at 37°C. The dissociated neurons were plated onto pre-coated PDL and growth factor-reduced matrigel (Cat# 354234, BD Biosciences, Franklin Lanes, NJ, USA) at a density of 90'000 cells per 14-mm coverslip. DRG neurons were cultured for 21 days. Until the first pulse with 5-fluoro-2'-deoxyuridine (FUdr), cells were cultured in c-DMEM in the presence of 100 ng/ml NGF (nerve growth factor) (Thermo Fisher Scientific, Rockford, IL, USA). To eliminate proliferating contaminant cells, cultures were pulsed twice for 48 hours with 20 µM FUdr (Sigma, St Louis, MO, USA), at 1 and 6 DIV. After the first FUdr pulse, the medium was changed to DMEM-Glutamax supplemented with 1 % B27 (Invitrogen, Thermo Fisher Scientific, Rockford, IL, USA), 1 % penicillin / streptomycin and 100 ng/ml NGF. After 21 days the medium was changed to 1x SATO diluted in DMEM F-12 supplemented with 1% B27, 100 ng/ml NGF, 5 ng/ml N-acetyl-L-cysteine, 5 mg/ml insulin (Cat# I9278, Sigma, St Louis, MO, USA) and 10 ng/ml D-biotin. At this stage, sorted oligodendrocyte progenitor cells were seeded onto the neurons at a density of 8’000 cells per coverslip. Infected oligodendrocyte progenitors were isolated by cell sorting. 48-hour post-infection, cells were detached (0.05 % Trypsin-EDTA), centrifuged, resuspended in FACS sorting buffer (1 mM EDTA, 25 mM HEPES, 2% FBS, 1x PBS (Ca2+/Mg+ free)) and passed through a cell strainer. The cell suspension was sorted for GFP in a FACSAria III (BD Biosciences, Allschwil, Switzerland) cell sorter. Co-cultures were maintained for 18 days *in vitro* and media replaced every 2 to 3 days.

*Quantitative RT-PCR*

Total RNA was extracted from oligodendrocyte cultures at 0, 3 and 6 days of differentiation (DOD 0, DOD 3 and DOD 6; where DOD 0 corresponds to the addition of 0.5 % FBS-supplemented SATO medium to the cultures; see oligodendrocyte precursor cells protocol). RNA was isolated using the Quick-RNA MicroPrep kit (ZymoResearch, Irvine, CA, USA) following the manufacturer’s recommendations. cDNA was synthesized from 0.5-2 μg total RNA using the SuperScript III reverse transcriptase (Thermo Fisher Scientific, Rockford, IL, USA). Quantitative RT-PCR was performed using iQ SYBR green supermix (Biorad, Hercules, CA, USA) in a iCycler iQ5 real-time PCR machine (Biorad, Hercules, CA, USA). The following program was used: 3 minutes at 94 °C, 40 steps of 15 seconds at 94 °C, 20 seconds at 60 °C, and 15 seconds at 72 °C. Used primers were: forward *Fscn1* 5’-AAAGATGAGCTCTTCGCCCT-3’, reverse *Fscn1* 5’-GATTGGCTGACAGGCCATT-3’, forward *Ywhaz* 5’-GATGAAGCCATTGCTGAACTTG-3’, reverse *Ywhaz* 5’-GTCTCCTTGGGTATCCGATGTC-3’ (Nelissen *et al.*, 2010).

*Immunostaining*

Immunostaining was performed to: (i) examine FSCN1 expression; (ii) analyze oligodendrocyte morphology following FSCN1 knockdown; and (iii) determine formation and length on internodes in myelinating co-cultures following FSCN1 knockdown. Cells were fixed in microtubule protection buffer (MP-PFA, 65 mM PIPES, 25 mM HEPES, 10 mM EGTA, 3 mM $MgCl_2$, pH 6.9 in 4 % paraformaldehyde) for 15 minutes at room temperature at DOD 6 (i), or 0, 3, and 6 (ii). After permeabilization in 0.1 % Triton X-100, 1x PBS, blocking was performed for 1 hour in 10 % normal goat serum, 1x PBS (blocking solution). Cells were incubated with primary antibodies diluted in blocking solution overnight at 4 °C, followed by secondary antibodies at room temperature for 1 hour.

For myelinating co-culture immunostaining, cells were fixed in MP-PFA buffer for 15 minutes at room temperature, followed by permeabilization and blocking (in 1 % HEPES buffer, 2.5 % Triton X-100, 1 % NGS and 1 % normal horse serum in 1x PBS) for 30 minutes at room temperature. Cells were incubated with primary antibodies diluted in the permeabilization/blocking solution for 1 hour at room temperature, followed by incubation with secondary antibodies for 1 hour at room temperature. After washing coverslips were mounted on Superfrost glass slides (Thermo Fisher Scientific, Rockford, IL, USA) with Fluoroshield mounting medium (Sigma, St Louis, MO, USA).

#### *Image acquisition and quantification*

Images were acquired onto a motorized inverted DMI 6000B epifluorescence microscope (Leica, Wetzlar, Germany) equipped with a digital ORCA-Flash4.0 vs2 Digital CMOS camera (C11440-22CU, Hamamatsu Photonics, Japan), with the following objectives: HC FL PLAN 10x / 0.25 dry (Cat# 11506307, Leica, Wetzlar, Germany); HC PL FLUOTAR L 20x / 0.40 dry (Cat# 11506243, Leica, Wetzlar, Germany); HC PL FLUOTAR L 40x / 0.60 dry (Cat# 11506203, Leica, Wetzlar, Germany).

The Internode formation index (= number of internodes per cell normalized to the mean of the control condition set to 100) index was calculated in two independent experiment with 9 to 16 cells analyzed per condition and experiment.

Olig2+ cells were classified according to their morphological complexity into 4 stages as previously described (Baer *et al.*, 2009): stage I mono/bipolar cells; stage II multipolar, primary branched cells; stage III multipolar, secondary or higher grade branched cells, and stage IV: branched cells with membrane enlargements. 3 independent experiments per time point (0, 3 and 6 DOD) were quantified, with at least 35 transduced oligodendrocytes (GFP+ Olig2+) quantified per experiment and time point.

#### *Experimental design*

Littermate and aged matched mice were randomly assigned to groups. No statistical method was used to predetermine sample size, but our sample sizes are similar to those generally employed in the field. All quantifications were done blindly by third party concealment of treatments.

#### *Statistical analysis*

Statistics were analyzed using GraphPad Prism vs6.01. Data were assumed to be normally distributed, but not formally tested. Variance was assumed to be equal between groups of data. Statistical significance was determined using an unpaired two sample Student's t-test for two group comparisons, while multiple group analysis was performed with one- or two-way analysis of variance (ANOVA) and post-hoc test as detailed in text and figures. Data show mean ± SEM of 3 independently run experiments unless otherwise specified in text and figures. Significance was set at $p < 0.05$ *, $p < 0.01$ **, $p < 0.001$ ***.

Supplementary Figure 1.

A

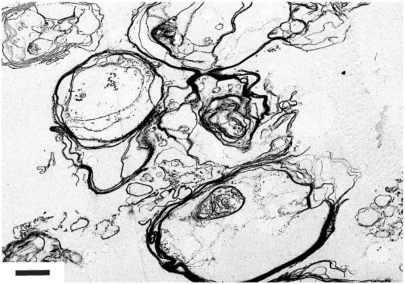

**Supplementary Figure 1. Myelin membranes isolation.** Representative micrograph of electron transmission microscope analysis of the pelleted myelin fraction showing enrichment in myelin membranes. Scale bar: 1 μm.

Supplementary Figure 2.

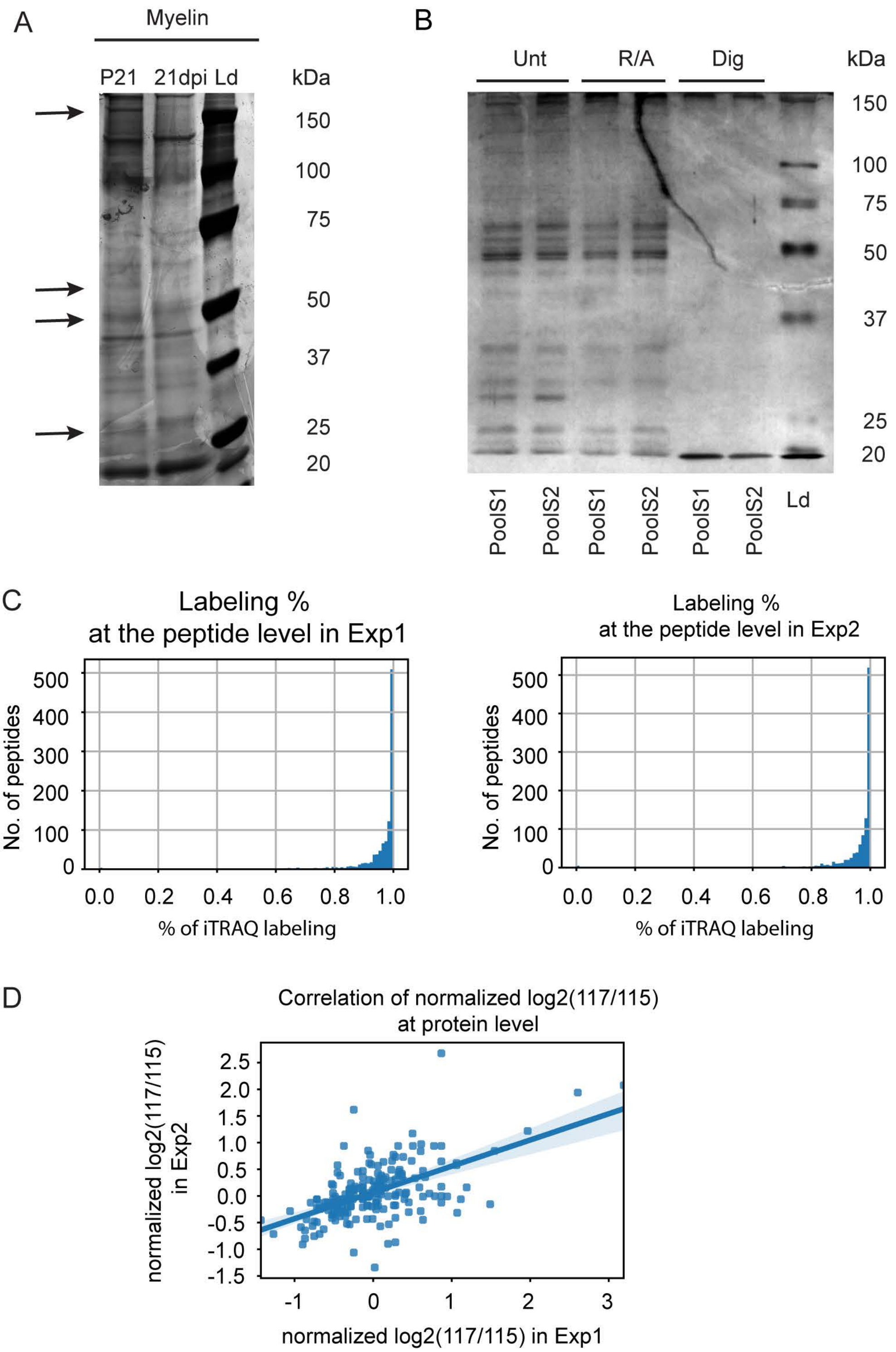


**Supplementary Figure 2. Myelin proteome deregulation following demyelination. (A)** Silver staining of protein lysates obtained from isolated myelin fractions (from the spinal cord of C57Bl/6 naïve P21 mice and mice that underwent a focal demyelinating lesion at 21 dpi) resolved on a 4-12% gradient Nu-PAGE. Differently expressed proteins (bands) are highlighted (arrows). **(B)** Silver staining of protein lysates of n = 2 two independent biological replicates for 21 dpi (each replicate obtained from 6-8 pooled spinal cord fragments, each 1 cm long around the lesion site) before and following reduction/alkylation, and subsequent trypsin digestion for iTRAQ labelling. **(C)** Histograms of the percentage iTRAQ labeling efficiency for the two iTRAQ experiments. **(D)** Correlation of normalized log2(117/115) values at the protein level across the two iTRAQ measurements (Pearson's correlation coefficient: 0.571911). Ld = Ladder, Unt = Untreated, R/A = Reduced and alkylated, Dig = Digested with Trypsin, Exp1 = experiment 1, Exp2 = experiment 2, PoolS1 = pooled samples 1, PoolS2 = pooled samples 2.

Supplementary Figure 3.

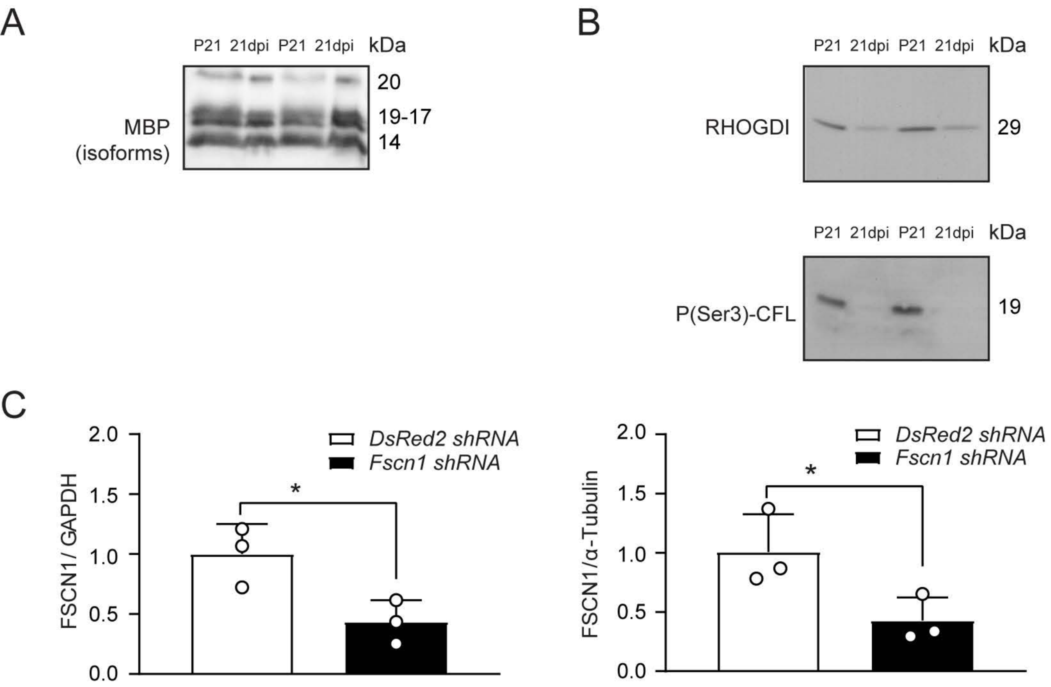


**Supplementary Figure 3. Confirmation of MBP, RhoGDI and cofilin expression levels in the remyelination proteome, and of knockdown of FCSCN1 by shRNA in oligodendrocytes in vitro. (A)** Immunoblot for MBP, showing similar expression in the remyelination versus the myelin naive proteome. **(B)** Immunoblots for RhoGDI and P(Ser3)-CFL showing lower expression in the remyelination versus the myelin naïve proteome. In (A) and (B) each sample from 8 pooled spinal cord fragments (each from a separate mouse) at P21 for naive myelin and at 21dpi from myelin formed by aOPCs after lysolecithin-induced demyelination. n = 2 independent set of samples / mice for each time point and condition. **(C)** Quantification graphs for FSCN1 immunoblot shown in Figure 3D, normalized to GAPDH and $\alpha$-Tubulin. n = 3 blots, each data point represents an independent experiment with an independent set of lysates from control (dsRed2 shRNA) and FSCN1 knockdown (FSCN1 shRNA) in cultured oligodendrocytes (unpaired one-tailed two sample Student's t test, FSCN1/GAPDH $P = 0.0169$, $t = 3.170$, FSCN1/$\alpha$-Tubulin $P = 0.0273$, $t = 2.690$; *, $P < 0.05$).

Supplementary Figure 4.

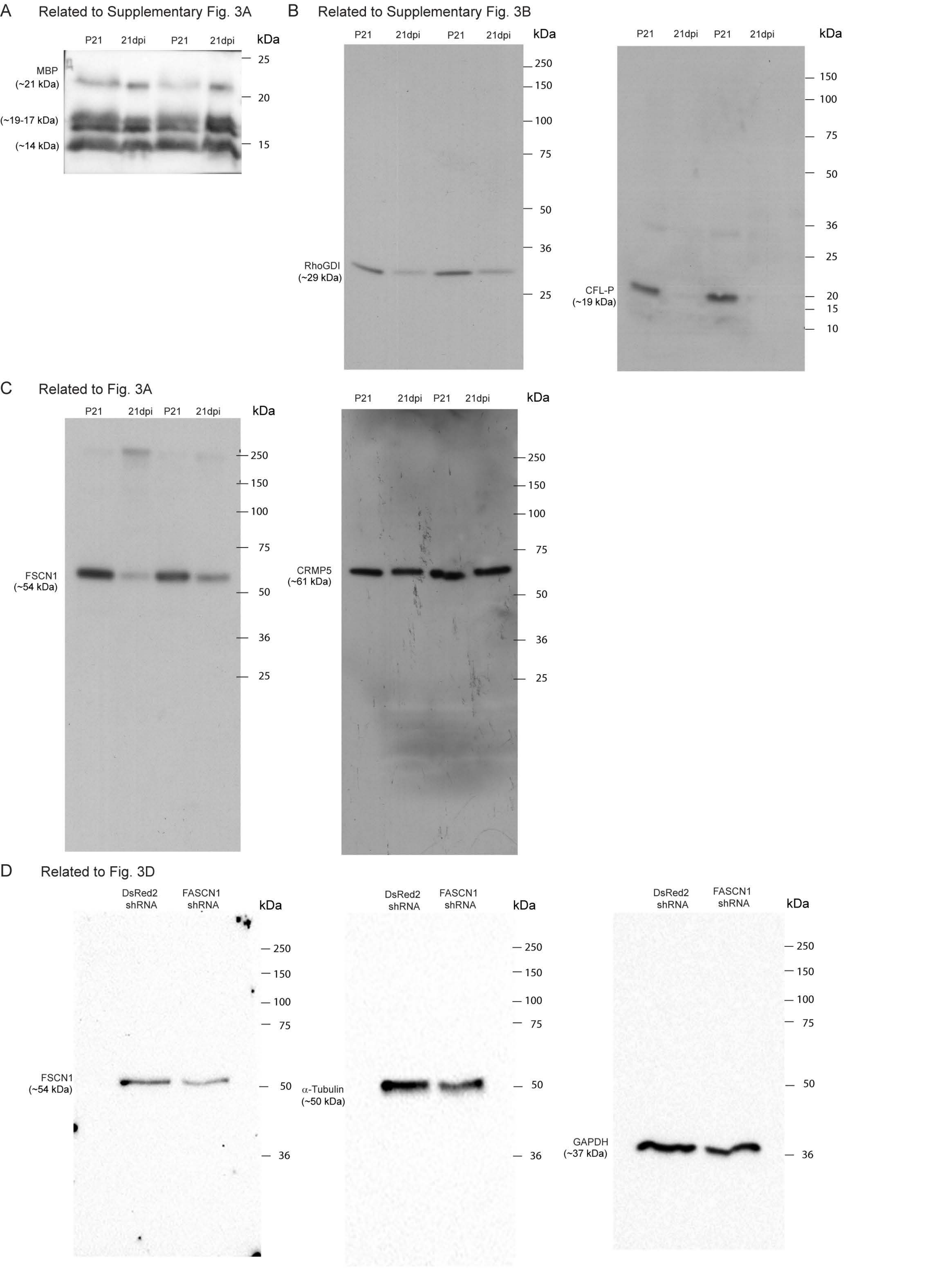


**Supplementary Figure 4. Full-length immunoblots.** Full-length immunoblots related to (**A, B**) Supplementary Fig. 3A, B, (**C**) Fig. 3A and (**D**) Fig. 3D. Gels were used as 10% (RhoGDI in B, FSCN1 and CRMP5 in C), 12% (FSCN1, Tubulin and GAPDH in D), and 14% (CFL-P in B) SDS-PAGE gels. After transfer, membrane in A was cut at 25 kD; Membrane in D was probed for FSCN1, stripped and re-probed for tubulin. Numbers refer to molecular weight size markers (kDa).

Supplementary Figure 5.

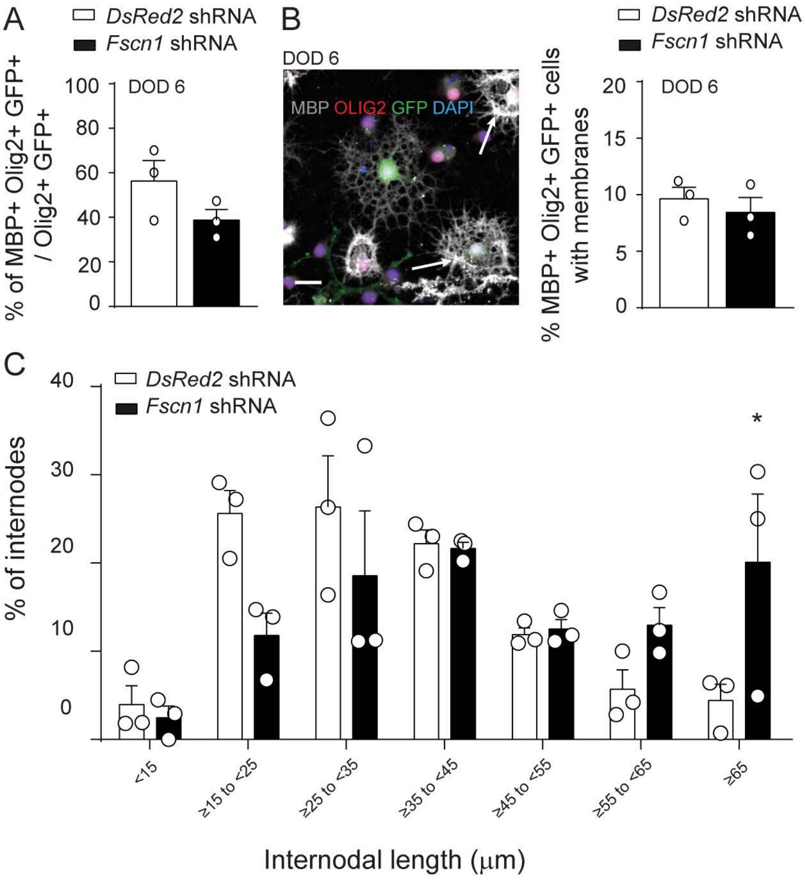


**Supplementary Figure 5. Oligodendrocyte maturation and internodal length are not majorly dependent upon FSCN1 expression. (A)** Bar graph showing the percentage of Olig2+ GFP+ cells expressing MBP as detected by immunocytochemistry after 6 DOD of OPCs *in vitro*, from control (*DsRed2* shRNA) and *Fscn1* shRNA transduced cells. *n* = 3 experiments (unpaired two-tailed two sample Student's t-test, *p* = 0.1695, *t* = 1.674). **(B)** Representative image of transfected (GFP+) fully differentiated (MBP+) oligodendrocytes (olig2+) forming membranes (arrows) and corresponding bar graph depicting the percentage of MBP+ Olig2+ GFP+ cells which formed membranes after 6 DOD *in vitro*, from control (*DsRed2* shRNA) and *Fscn1* shRNA transduced cells. *n* = 3 experiments (unpaired two-tailed two sample Student's t-test, *p* = 0.5121, *t* = 0.7186). Scale bar 20 μm. **(C)** Bar graph showing the percentage of internodes assigned to binning of internodal length distribution from control (*DsRed2* shRNA) and *Fscn1* shRNA transduced oligodendrocytes, GFP+ sorted and induced to myelinate axons ex vivo after 18 days, *n* = 3 experiments (2-way Anova with Sidak's multiple comparisons test, *Fscn1* shRNA versus *DsRed2* shRNA <15 μm *p* > 0.9999, 15 to 25 μm *p* = 0.0796, 25 to 35 μm *p* = 0.6486, 35 to 45 μm *p* > 0.9999, 45 to 55 μm *p* > 0.9999, 55 to 65 μm *p* = 0.7219, > 65 μm *p* = 0.0337), * p < 0.05. Bars represent mean ± SEM. DOD = days of differentiation.

**Supplementary Table 1. The remyelination proteome.** List of proteins showing differential expression in the remyelination proteome compared to the developmental myelin proteome (21 dpi compared to young adult myelin at P21). Only statistically significant changes (applying a threshold of ≥ 2 unique peptides in each of the two biological replicates, ≥ 0.9 protein probability score in each of the two biological replicates, and ≥ 1.2 mean fold change as either up- or down-regulation in both experiments) are shown.

| ENTREZ gene_ID | UP_SYMBOL | Description | Protein Prob Rep 1 | # unique peptides Rep 1 | Protein Prob Rep 2 | # unique peptides Rep 2 | Mean Ratio | Error |
|---|---|---|---|---|---|---|---|---|
| 13627 | EF1A1_MOUSE | Elongation factor 1-alpha 1 | 1 | 9 | 1 | 2 | 0.30 | 0.08 |
| 15926 | IDHC_MOUSE | Isocitrate dehydrogenase [NADP] cytoplasmic | 1 | 4 | 1 | 2 | 0.32 | 0.09 |
| 64383 | SIRT2_MOUSE | NAD-dependent deacetylase sirtuin-2 | 1 | 5 | 1 | 8 | 0.34 | 0.15 |
| 12317 | CALR_MOUSE | Calreticulin | 1 | 4 | 1 | 2 | 0.35 | 0.08 |
| 17136 | MAG_MOUSE | Myelin-associated glycoprotein | 1 | 4 | 1 | 5 | 0.35 | 0.19 |
| 70350 | BASP1_MOUSE | Brain acid soluble protein 1 | 1 | 2 | 1 | 4 | 0.35 | 0.09 |
| 22142 | TBA1A_MOUSE | Tubulin alpha-1A chain | 1 | 9 | 1 | 5 | 0.36 | 0.01 |
| 22627 | 1433E_MOUSE | 14-3-3 protein epsilon | 1 | 6 | 1 | 3 | 0.38 | 0.04 |
| 11947 | ATPB_MOUSE | ATP synthase subunit beta, mitochondrial | 1 | 8 | 1 | 8 | 0.39 | 0.08 |
| 14827 | PDIA3_MOUSE | Protein disulfide-isomerase A3 | 1 | 3 | 1 | 5 | 0.40 | 0.09 |
| 109905 | RAP1A_MOUSE | Ras-related protein Rap-1b | 1 | 2 | 1 | 2 | 0.40 | 0.08 |
| 78294 | UBIQ_MOUSE | Ubiquitin | 1 | 3 | 1 | 4 | 0.40 | 0.05 |
| 12799 | CN37_MOUSE | 2',3'-cyclic-nucleotide 3'-phosphodiesterase | 1 | 32 | 1 | 37 | 0.40 | 0.04 |
| 21779 | TRFE_MOUSE | Serotransferrin | 1 | 6 | 1 | 3 | 0.40 | 0.06 |
| 14086 | FSCN1_MOUSE | Fascin | 1 | 2 | 1 | 2 | 0.40 | 0.17 |
| 232975 | AT1A3_MOUSE | Sodium/potassium-transporting ATPase subunit alpha-3 | 1 | 14 | 1 | 19 | 0.41 | 0.10 |
| 26950 | VISL1_MOUSE | Visinin-like protein 1 | 1 | 4 | 1 | 4 | 0.41 | 0.03 |

| ENTREZ gene_ID | UP_SYMBOL | Description | Protein Prob Rep 1 | # unique peptides Rep 1 | Protein Prob Rep 2 | # unique peptides Rep 2 | Mean Ratio | Error |
|---|---|---|---|---|---|---|---|---|
| 67300 | CLH_MOUSE | Clathrin heavy chain 1 | 1 | 8 | 1 | 7 | 0.42 | 0.04 |
| 11931 | AT1B1_MOUSE | Sodium/potassium-transporting ATPase subunit beta-1 | 1 | 5 | 1 | 6 | 0.42 | 0.11 |
| 14681 | GNAO_MOUSE | Guanine nucleotide-binding protein G(o) subunit alpha | 1 | 4 | 1 | 2 | 0.42 | 0.08 |
| 12631 | COF1_MOUSE | Cofilin-1 | 1 | 10 | 1 | 7 | 0.43 | 0.04 |
| 22240 | DPYL3_MOUSE | Dihydropyrimidinase-related protein 3 | 1 | 12 | 1 | 6 | 0.43 | 0.04 |
| 12716 | KCRU_MOUSE | Creatine kinase U-type, mitochondrial | 1 | 2 | 1 | 2 | 0.44 | 0.17 |
| 22195 | UB2L3_MOUSE | Ubiquitin-conjugating enzyme E2 L3 | 1 | 2 | 1 | 3 | 0.44 | 0.16 |
| 276770 | IF5A1_MOUSE | Eukaryotic translation initiation factor 5A-1 | 1 | 3 | 1 | 2 | 0.45 | 0.04 |
| 18477 | PRDX1_MOUSE | Peroxiredoxin-1 | 1 | 8 | 1 | 8 | 0.45 | 0.05 |
| 0 | SPTA2_MOUSE | Spectrin alpha chain, brain | 1 | 5 | 1 | 6 | 0.45 | 0.08 |
| 268373 | PPIA_MOUSE | Peptidyl-prolyl cis-trans isomerase A | 1 | 11 | 1 | 12 | 0.45 | 0.05 |
| 15516 | HS90B_MOUSE | Heat shock protein HSP 90-beta | 1 | 6 | 1 | 12 | 0.46 | 0.05 |
| 21881 | TKT_MOUSE | Transketolase | 1 | 6 | 1 | 5 | 0.46 | 0.06 |
| 192662 | GDIR1_MOUSE | Rho GDP-dissociation inhibitor 1 | 1 | 2 | 1 | 3 | 0.47 | 0.10 |
| 17319 | MIF_MOUSE | Macrophage migration inhibitory factor | 1 | 4 | 1 | 4 | 0.47 | 0.09 |
| 11674 | ALDOA_MOUSE | Fructose-bisphosphate aldolase A | 1 | 16 | 1 | 18 | 0.48 | 0.04 |
| 15510 | CH60_MOUSE | 60 kDa heat shock protein, mitochondrial | 1 | 5 | 1 | 4 | 0.49 | 0.05 |
| 12140 | FABP7_MOUSE | Fatty acid-binding protein, brain | 1 | 2 | 1 | 3 | 0.49 | 0.11 |

| ENTREZ gene_ID | UP_SYMBOL | Description | Protein Prob Rep 1 | # unique peptides Rep 1 | Protein Prob Rep 2 | # unique peptides Rep 2 | Mean Ratio | Error |
|---|---|---|---|---|---|---|---|---|
| 269523 | TERA_MOUSE | Transitional endoplasmic reticulum ATPase | 1 | 6 | 1 | 10 | 0.49 | 0.05 |
| 21672 | PRDX2_MOUSE | Peroxiredoxin-2 | 1 | 5 | 1 | 5 | 0.49 | 0.06 |
| 22628 | 1433G_MOUSE | 14-3-3 protein gamma | 1 | 6 | 1 | 6 | 0.50 | 0.03 |
| 11461 | ACTB_MOUSE | Actin, cytoplasmic 2 | 1 | 12 | 1 | 7 | 0.50 | 0.04 |
| 12934 | DPYL2_MOUSE | Dihydropyrimidinase-related protein 2 | 1 | 24 | 1 | 22 | 0.50 | 0.05 |
| 15481 | HSP7C_MOUSE | Heat shock cognate 71 kDa protein | 1 | 26 | 1 | 15 | 0.51 | 0.03 |
| 11749 | ANXA6_MOUSE | Annexin A6 | 1 | 5 | 1 | 5 | 0.51 | 0.04 |
| 16765 | STMN1_MOUSE | Stathmin-2 | 1 | 2 | 1 | 5 | 0.51 | 0.03 |
| 72948 | TPPP_MOUSE | Tubulin polymerization-promoting protein | 1 | 2 | 1 | 2 | 0.51 | 0.10 |
| 14719 | AATM_MOUSE | Aspartate aminotransferase, mitochondrial | 1 | 3 | 1 | 4 | 0.51 | 0.11 |
| 13628 | EF1A2_MOUSE | Elongation factor 1-alpha 2 | 1 | 2 | 1 | 5 | 0.52 | 0.05 |
| 11964 | VATA_MOUSE | V-type proton ATPase catalytic subunit A | 1 | 5 | 1 | 4 | 0.52 | 0.10 |
| 17433 | MOBP_MOUSE | Myelin-associated oligodendrocyte basic protein | 1 | 2 | 1 | 3 | 0.52 | 0.11 |
| 22153 | TBB4_MOUSE | Tubulin beta-4 chain | 1 | 8 | 1 | 8 | 0.52 | 0.08 |
| 51792 | 2AAA_MOUSE | Serine/threonine-protein phosphatase 2A 65 kDa regulatory subunit A alpha isoform | 1 | 7 | 1 | 4 | 0.52 | 0.11 |
| 22201 | UBA1_MOUSE | Ubiquitin-like modifier-activating enzyme 1 | 1 | 7 | 1 | 4 | 0.52 | 0.07 |
| 320981 | ENPP6_MOUSE | Ectonucleotide pyrophosphatase/phosphodiesterase family member 6 | 1 | 2 | 1 | 2 | 0.52 | 0.08 |

| ENTREZ gene_ID | UP_SYMBOL | Description | Protein Prob Rep 1 | # unique peptides Rep 1 | Protein Prob Rep 2 | # unique peptides Rep 2 | Mean Ratio | Error |
|---|---|---|---|---|---|---|---|---|
| 16832 | LDHB_MOUSE | L-lactate dehydrogenase B chain | 1 | 5 | 1 | 7 | 0.52 | 0.05 |
| 15519 | HS90A_MOUSE | Heat shock protein HSP 90-alpha | 1 | 16 | 1 | 14 | 0.53 | 0.06 |
| 11973 | VATE1_MOUSE | V-type proton ATPase subunit E 1 | 1 | 5 | 1 | 3 | 0.53 | 0.09 |
| 0 | MAP2_MOUSE | Microtubule-associated protein 2 | 1 | 2 | 1 | 5 | 0.53 | 0.08 |
| 17441 | MOG_MOUSE | Myelin-oligodendrocyte glycoprotein | 1 | 4 | 1 | 3 | 0.53 | 0.12 |
| 13424 | DYHC1_MOUSE | Cytoplasmic dynein 1 heavy chain 1 | 1 | 6 | 1 | 4 | 0.53 | 0.11 |
| 17967 | NCAM1_MOUSE | Neural cell adhesion molecule 1 | 1 | 2 | 1 | 3 | 0.54 | 0.12 |
| 14567 | GDIA_MOUSE | Rab GDP dissociation inhibitor alpha | 1 | 6 | 1 | 2 | 0.55 | 0.11 |
| 17448 | MDHM_MOUSE | Malate dehydrogenase, mitochondrial | 1 | 8 | 1 | 8 | 0.55 | 0.04 |
| 20910 | STXB1_MOUSE | Syntaxin-binding protein 1 | 1 | 4 | 1 | 4 | 0.55 | 0.15 |
| 22631 | 1433Z_MOUSE | 14-3-3 protein zeta/delta | 1 | 8 | 1 | 7 | 0.55 | 0.04 |
| 56370 | TAGL3_MOUSE | Transgelin-3 | 0.9999 | 2 | 1 | 2 | 0.57 | 0.06 |
| 11429 | ACON_MOUSE | Aconitate hydratase, mitochondrial | 1 | 9 | 1 | 7 | 0.57 | 0.05 |
| 11676 | ALDOC_MOUSE | Fructose-bisphosphate aldolase C | 1 | 12 | 1 | 11 | 0.58 | 0.04 |
| 18648 | PGAM1_MOUSE | Phosphoglycerate mutase 1 | 1 | 10 | 1 | 11 | 0.58 | 0.04 |
| 18655 | PGK1_MOUSE | Phosphoglycerate kinase 1 | 1 | 12 | 1 | 10 | 0.58 | 0.04 |
| 22154 | TBB5_MOUSE | Tubulin beta-5 chain | 1 | 4 | 1 | 5 | 0.59 | 0.08 |
| 54683 | PRDX5_MOUSE | Peroxiredoxin-5, mitochondrial | 1 | 2 | 1 | 5 | 0.59 | 0.21 |
| 15528 | CH10_MOUSE | 10 kDa heat shock protein, mitochondrial | 1 | 3 | 1 | 3 | 0.59 | 0.06 |
| 18475 | PA1B2_MOUSE | Platelet-activating factor acetylhydrolase IB subunit beta | 1 | 2 | 1 | 2 | 0.60 | 0.11 |
| 11464 | ACTC_MOUSE | Actin, alpha skeletal muscle | 1 | 4 | 1 | 3 | 0.60 | 0.01 |
| 13806 | ENOA_MOUSE | Alpha-enolase | 1 | 14 | 1 | 13 | 0.61 | 0.03 |
| 0 | G6PI_MOUSE | Glucose-6-phosphate isomerase | 1 | 5 | 1 | 3 | 0.61 | 0.05 |

| ENTREZ gene_ID | UP_SYMBOL | Description | Protein Prob Rep 1 | # unique peptides Rep 1 | Protein Prob Rep 2 | # unique peptides Rep 2 | Mean Ratio | Error |
|---|---|---|---|---|---|---|---|---|
| 109801 | LGUL_MOUSE | Lactoylglutathione lyase | 1 | 2 | 1 | 3 | 0.62 | 0.03 |
| 18102 | NDKA_MOUSE | Nucleoside diphosphate kinase A | 1 | 5 | 1 | 8 | 0.62 | 0.08 |
| 18039 | NFL_MOUSE | Neurofilament light polypeptide | 1 | 16 | 1 | 16 | 0.62 | 0.04 |
| 14433 | G3P_MOUSE | Glyceraldehyde-3-phosphate dehydrogenase | 1 | 15 | 1 | 15 | 0.63 | 0.03 |
| 268860 | GABT_MOUSE | 4-aminobutyrate aminotransferase, mitochondrial | 1 | 3 | 1 | 2 | 0.63 | 0.15 |
| 18746 | KPYM_MOUSE | Pyruvate kinase isozymes M1/M2 | 1 | 15 | 1 | 15 | 0.64 | 0.04 |
| 11636 | KAD1_MOUSE | Adenylate kinase isoenzyme 1 | 1 | 2 | 1 | 3 | 0.64 | 0.06 |
| 380684 | NFH_MOUSE | Neurofilament heavy polypeptide | 1 | 3 | 1 | 6 | 0.64 | 0.04 |
| 14718 | AATC_MOUSE | Aspartate aminotransferase, cytoplasmic | 1 | 6 | 1 | 3 | 0.64 | 0.11 |
| 23980 | PEBP1_MOUSE | Phosphatidylethanolamine-binding protein 1 | 1 | 4 | 1 | 3 | 0.64 | 0.11 |
| 22151 | TBB2A_MOUSE | Tubulin beta-2A chain | 1 | 7 | 1 | 7 | 0.66 | 0.04 |
| 13807 | ENOG_MOUSE | Gamma-enolase | 1 | 12 | 1 | 9 | 0.66 | 0.05 |
| 14862 | GSTM1_MOUSE | Glutathione S-transferase Mu 1 | 1 | 6 | 1 | 7 | 0.67 | 0.07 |
| 14645 | GLNA_MOUSE | Glutamine synthetase | 1 | 5 | 1 | 6 | 0.68 | 0.09 |
| 12889 | CPLX1_MOUSE | Complexin-1 | 1 | 4 | 1 | 4 | 0.69 | 0.03 |
| 14661 | DHE3_MOUSE | Glutamate dehydrogenase 1, mitochondrial | 1 | 2 | 1 | 3 | 0.69 | 0.14 |
| 11747 | ANXA5_MOUSE | Annexin A5 | 1 | 7 | 1 | 7 | 0.70 | 0.06 |
| 18103 | NDKB_MOUSE | Nucleoside diphosphate kinase B | 1 | 5 | 1 | 3 | 0.70 | 0.06 |
| 21991 | TPIS_MOUSE | Triosephosphate isomerase | 1 | 7 | 1 | 9 | 0.71 | 0.02 |
| 226180 | AINX_MOUSE | Alpha-internexin | 1 | 7 | 1 | 3 | 0.71 | 0.05 |
| 17449 | MDHC_MOUSE | Malate dehydrogenase, cytoplasmic | 1 | 5 | 1 | 7 | 0.71 | 0.04 |
| 22152 | TBB3_MOUSE | Tubulin beta-3 chain | 1 | 11 | 1 | 11 | 0.71 | 0.04 |
| 12308 | CALB2_MOUSE | Calretinin | 1 | 4 | 1 | 4 | 0.74 | 0.09 |

| ENTREZ gene_ID | UP_SYMBOL | Description | Protein Prob Rep 1 | # unique peptides Rep 1 | Protein Prob Rep 2 | # unique peptides Rep 2 | Mean Ratio | Error |
|---|---|---|---|---|---|---|---|---|
| 20655 | SODC_MOUSE | Superoxide dismutase [Cu-Zn] | 1 | 4 | 1 | 4 | 0.79 | 0.03 |
| 12709 | KCRB_MOUSE | Creatine kinase B-type | 1 | 21 | 1 | 21 | 0.80 | 0.03 |
| 11807 | APOA2_MOUSE | Apolipoprotein A-II | 1 | 2 | 1 | 2 | 1.68 | 0.34 |
| 15121 | HBA_MOUSE | Hemoglobin subunit alpha | 1 | 10 | 1 | 9 | 2.67 | 0.04 |
| 15129 | HBB1_MOUSE | Hemoglobin subunit beta-1 | 1 | 17 | 1 | 10 | 3.60 | 0.05 |
